# Detecting local changes in chromatin architecture with false discovery control

**DOI:** 10.1101/2020.09.03.281972

**Authors:** Hillary Koch, Tao Yang, Maxim Imakaev, Ross C. Hardison, Qunhua Li

**Affiliations:** Department of Statistics, Pennsylvania State University; Regeneron Pharmaceuticals, Inc; Institute for Medical Engineering and Science, Massachusetts Institute of Technology; Department of Biochemistry and Molecular Biology, Pennsylvania State University

## Abstract

Hi-C experiments are a powerful means to describe the organization of chromatin interactions genome-wide. By using Hi-C data to identify differentially organized genomic regions, relationships between this organization, gene expression, and cell identity may be established. However, Hi-C data exhibit a unique and challenging spatial structure, as genomic loci can show strong correlations when they are nearby in 3D space within the nucleus or 1D space along the chromosome. Consequently, the development of methods that can accurately detect differences between Hi-C samples while controlling false discoveries has remained difficult. To meet this need, we introduce a spatial modeling approach based on sliding window statistics. Using polymer simulations, we illustrate the improved power and precision of our method to identify differentially interacting genomic regions. We further demonstrate our method’s ability to reveal biologically meaningful changes in chromatin architecture through two data analyses concerning the loss of architectural and chromatin remodeling proteins.

## Introduction

High-throughput methods that probe 3D genome architecture *in situ*, Hi-C in particular^1^, have provided new means to study factors affecting transcriptional control within the cell genome-wide. The increasing availability of Hi-C data makes it possible to study the role of genomic interactions in processes such as cell differentiation^2,3,4^ and pathogenesis^5,6,7^. For example, beyond somatic mutations and cancer genes, factors that affect genetic and epigenetic regulation have been shown to affect cancer evolution^8^. Specifically, the dysregulation of topologically-associated domains (TADs)^9^, insulated self-interacting genomic regions, has a demonstrated effect on gene expression in perturbed cells^10,11^. In order to understand how changes in the 3D interaction landscape impact cell function, one needs a way to correctly identify the structural differences in chromatin organization between cells sampled from different biological conditions.

To this end, several ad-hoc methods have been developed for the differential analysis of Hi-C contact matrices. One common methodological approach involves identifying differential peaks, or chromatin interaction hotspots, in Hi-C matrices^12,13,14,15^. However, these results are often irreproducible due to their susceptibility to high stochasticity inherent to Hi-C data^16^. Another approach involves calling TADs in two conditions, then comparing the called boundaries^13,17,18,19,20^. However, TAD callers identify boundaries inconsistently even within replicate experiments^21,22^, raising questions about the suitability of this approach for reliably identifying differences between experimental conditions.

Notably, many methods for identifying differences between Hi-C datasets suffer from at least one of two critical drawbacks. The first is an inability to statistically quantify confidence in the identified differences, and second is some amount of agnosticism to the most salient feature of Hi-C data–its complex spatial dependence. That is, Hi-C contact maps harbor both a 1D and 3D spatial dependence; not only are nearby regions along the genome correlated with one another, resulting in a distance dependence effect^23,24^, but a dependence induced by preferential within-TAD interactions results in blocks along the diagonal of the Hi-C contact map. Ignoring such evident dependencies in the data leads to compounding problems with sensitivity and false discovery control when calling significantly differential interactions^25,26,27^.

To address these critical concerns, we developed a principled statistical approach to identify differences between Hi-C contact maps, called locdiffr (analysis of LOCal DIFFerences in chromatin aRchitecture). Our method accounts for the well-known, complex sources of spatial dependence in Hi-C experiments while providing confidence estimates in differences at each location, permitting more accurate false discovery control. We compare locdiffr to existing tools using simulations of differing experimental conditions from polymer models, where the ground truth is known. Finally, we apply locdiffr to two Hi-C datasets, demonstrating its robustness and revealing biological insights into relationships between chromatin remodeling and changes in epigenetic states.

## Results

### Overview of approach to detecting regions of differential interactions

Our approach to identifying local changes in chromatin architecture employs two major steps:

1. Compute a score that evinces support for differential interactions within a sliding window across a given chromosome
2. Test for which windows show statistically significant support for containing differentially interacting genomic regions

We address preferential within-TAD interactions and the Hi-C distance dependence effect by using a scoring rule that accounts for these facets of Hi-C data in Step 1 (see next section). Moreover, we account for the correlation between scores computed within nearby windows through the utilization of a unified statistical framework for modeling and testing that accounts for spatial dependence in Step 2 (see *Modeling the SCC sliding window statistic*). Altogether, our approach cohesively addresses well-described types of spatial dependence evident in Hi-C data.

#### SCC sliding window statistic

Step 1 requires a scoring rule for differentially interacting genomic regions that accounts for preferential within-TAD interactions and the Hi-C distance dependence effect. Previously, Yang et al.^24^ introduced HiCRep, a tool to assess the reproducibility of Hi-C data. The authors demonstrated that the distance dependence effect, which manifests as higher contact frequencies near the diagonal of the Hi-C contact map, biases the commonly-used measures of correlation, such as Pearson correlation, between two samples. They therefore developed the stratum-adjusted correlation coefficient (SCC), which measures similarity between two Hi-C samples while accounting for the decay of signal with increasing distance from the Hi-C matrix diagonal.

Yang et al. established that the SCC is a useful tool for measuring agreement between two Hi-C experiments, and that it can even recapitulate known relationships among cell types when comparing several Hi-C samples. We therefore adapted this tool to measure *local* agreement between samples. By computing the SCC within a sliding window, instead of between two whole Hi-C contact matrices, we can obtain such a local measure of agreement between two Hi-C contact matrices. That is, for a fixed sliding window that spans *B*_*w*_ bins along the matrix diagonal, and two Hi-C experiments each conducted at a resolution yielding *B* bins, one can compute the SCC score between these two experiments in a total of *W* = *B − B*_*w*_ + 1 windows (see Fig. 1a). We use a triangular sliding window, which traverses the Hi-C matrix one bin at a time, to capture the hallmark TAD shape. Windows with a relatively high local SCC score correspond to agreement between samples, while those with low SCC scores correspond to windows showing a difference between samples. We term the set of SCC scores computed within these *W* windows as the SCC sliding window statistic.

**Figure 1:**
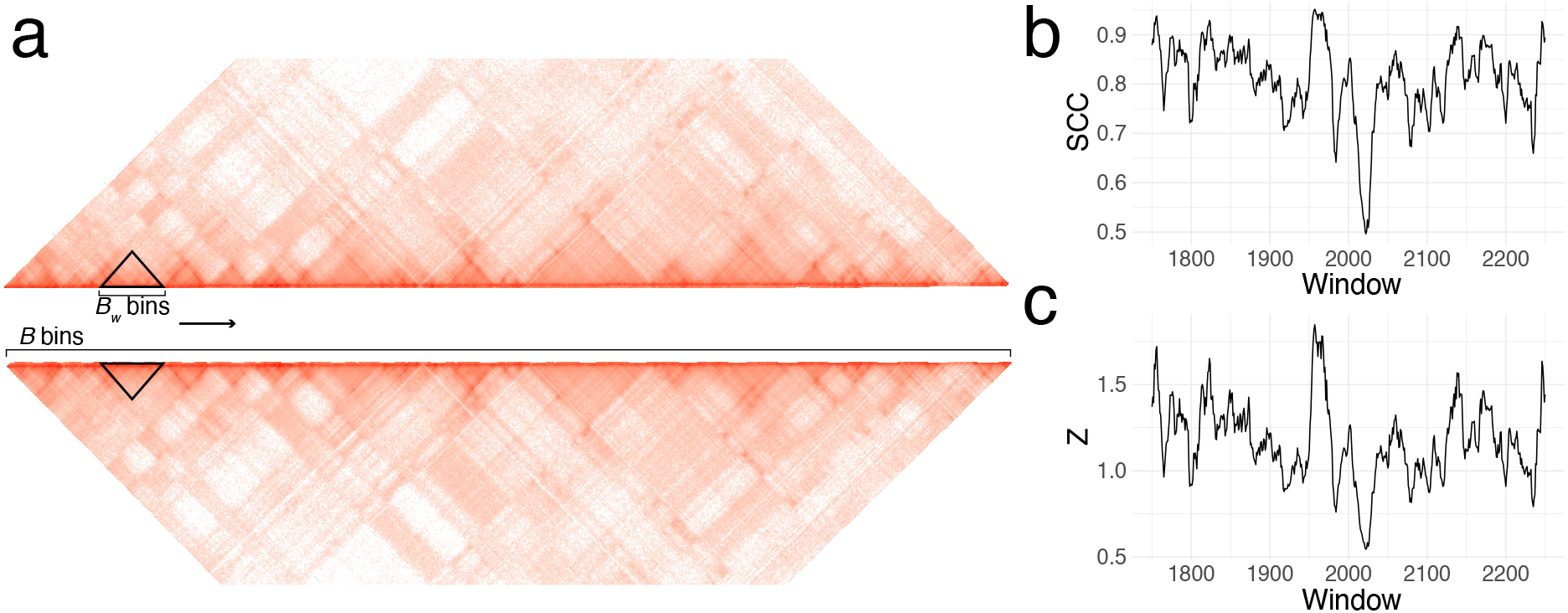
Diagram of locdiffr in a wild type vs. knockout experiment. **a**, The stratum-adjusted correlation coefficient (SCC) is computed in a sliding window between two Hi-C contact matrices from a region of chromosome 9 of control and Chd4-knockout murine cells^28^. **b**, The SCC scores for the shown scan and **c**, the SCC scores transformed into *z*-scores for the shown scan.

Describing local agreement between two Hi-C experiments with the SCC sliding window statistic accounts for the 3D distance dependence apparent in Hi-C data, yet correlation between nearby windows still remains. That is, the SCC scores from proximal windows are correlated because 1) the SCC scores are computed within a sliding window, and therefore two overlapping windows are likely to exhibit similar scores and 2) genomic regions that are close in 1D are likely to behave similarly. Therefore, in the next step, we explicitly model this behavior when testing for significantly differentially interacting genomic regions.

#### Modeling the SCC sliding window statistic as a stochastic process

In order to capture spatial dependence among SCC scores computed within proximal windows, we leverage Gaussian processes. Let *S ⊆ ℛ* be a spatial domain. Then, a stochastic process *Z*(***s***) indexed by spatial locations ***s*** *∈ S* is called a Gaussian process if, for any set of locations *{****s***_1_, *…*, ***s***_*w*_*}*, (*Z*(***s***_1_), *…, Z*(***s***_*w*_))has a multivariate normal distribution^29^. Gaussian processes have been widely applied to the modeling of time series^30,31^, spatial^32,33,34,35^, and spatio-temporal data to capture the expected dependencies among observations occurring nearby in time and space. Likewise, a series of dependent scores computed within a spatially-referenced sliding window could suitably be captured by a Gaussian process model.

As a correlation coefficient, the SCC score *ρ ∈* [*−*1, 1] is not Gaussian. By applying Fisher’s *z*-transformation element-wise^36^, we transform the computed SCC sliding window statistic into approximately standard normal random variables. That is, letting ***ρ*** be a *W*-vector of SCC scores computed within windows *s*_1_, *…, s*_*W*_,

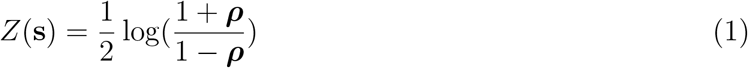

approximately follows a Gaussian process describing local agreement between two Hi-C matrices. Since equation (1) is a monotonic transformation of ***ρ***, higher values of *Z*(***s***) correspond to more local agreement, while lower values correspond to less local agreement between the samples.

We model this process as having a smooth, underlying background (i.e., the mean function *µ*) with an exponential covariance and iid noise (i.e., the covariance function *C*):

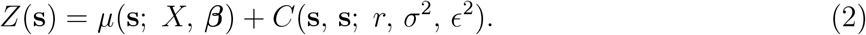

Here, *µ*(***s***; *X*, ***β***) := *µ*(***s***) = *X****β*** where *X* is a *W × p* cubic B-spline basis matrix on *S* and ***β*** is a *p*-vector of coefficients, *σ*^2^ is the marginal exponential variance or *partial sill, r* is the spatial range, and *σ*^2^*ϵ*^2^ is the iid error variance or *nugget*. The selected covariance function implies *weak stationarity* of the process, meaning that the covariance between any two locations in the process can be summarized by the distance between those locations. That is, for covariance function *C* and separation vector ***h***, we have *C*(***s, s*** + ***h***) = *C*(***h***)^37^; for the exponential covariance function in particular, we have *C*(*h*; *r, σ*^2^, *ϵ*^2^) := *C* = *σ*^2^*ϵ*^2^ + *σ*^2^ exp *{−h/r}*, such that the covariance between two spatial locations decays exponentially towards the sill (partial sill + nugget) as the separation between them grows.

Fitting the model in equation (2) is known to be computationally demanding when the dimension of the data is large, rendering an analysis with *∼* 10^3^ *−* 10^6^ spatial locations prohibitive or impossible^38^. To tackle this, Datta et al. (2016)^39^ introduced the nearest-neighbor Gaussian process (NNGP) for approximating Gaussian processes such as the one in equation (2). That is, letting

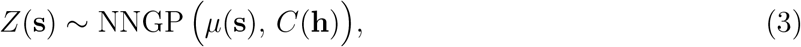

one assumes that the spatial dependence inherent to process *Z*(***s***) can be reasonably approximated by using only the *k*-nearest neighbors of each spatial location. By using this local information to approximate a standard Gaussian process, NNGPs are highly scalable as they circumvent the need to store and invert large covariance matrices during model fitting. For example, the Hi-C data for human chromosome 1 at 50 kilobase (kb) resolution contains around 5,000 bins. Therefore, comparing Hi-C matrices with the SCC sliding window statistic will produce data with a large number of spatial locations. This necessitates a more computationally efficient means of statistical modeling and inference, making approximation of the transformed SCC sliding window statistic by NNGPs a natural choice.

As will be discussed in the following section, we do inference for this model using Markov chain Monte Carlo (MCMC). Therefore, we cast our NNGP into the Bayesian framework, hierarchically representing the model as

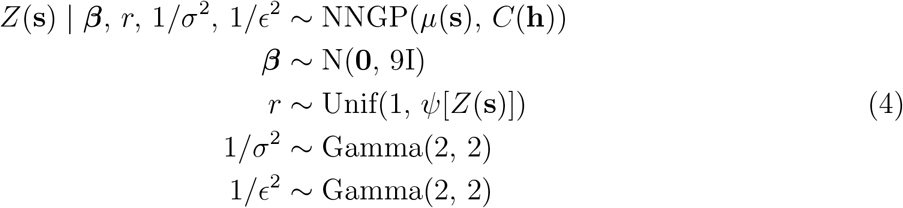

where *I* is the *p × p* identity matrix and *ψ* provides an upper bound on *r* by estimating the effective range of the process *Z*(***s***) (see Supplementary Section S1.1). This model benefits from increased experimental replicates, as each additional pair of replicates increases the sample size, resulting in more accurate parameter estimation (Supplementary equations (15) and (16)). The priors in (4) were chosen to be only lightly informative, given that we know that the SCC sliding window statistic is approximately *z*-distributed. We made this choice because wholely uninformative priors could make inference too difficult given the small sample sizes typical of Hi-C experiments.

#### Identifying differentially-interacting genomic regions with false discovery control

A significantly differentially-interacting genomic region i s one f or which we a re c onfident th at the transformed SCC score in window *s*_*w*_, *Z*(*s*_*w*_), is significantly less than the background in window *s*_*w*_, *µ*(*s*_*w*_). To measure our confidence that this is indeed the c ase at e ach spatial location *s* _*w*_, we test the hypothesis

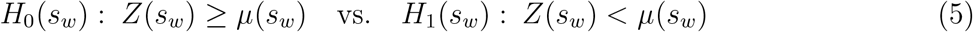

for *w ∈ {*1, *…, W}*. For a fixed window size, this results in *W* simultaneous hypothesis tests that are correlated. We can adjust for the multiple tests by applying false discovery controlling procedures^40,41,42^. First, define *θ*(*s*_*w*_) = 𝕝[*Z*(*s*_*w*_) *< µ*(*s*_*w*_)] as the indicator that the test for window *s*_*w*_ truly lies in the rejection region, and *δ*(*s*_*w*_) be the decision to accept (*δ*(*s*_*w*_) = 0) or reject (*δ*(*s*_*w*_) = 1) the null hypothesis for window *s*_*w*_. Then, given the set of *n* observed NNGPs ***Z*** = *{Z*_1_(***s***), *…, Z*_*n*_(***s***) *}*, the Bayesian false discovery rate^43,44^ (FDR) is the expected value of the false discovery proportion (FDP):

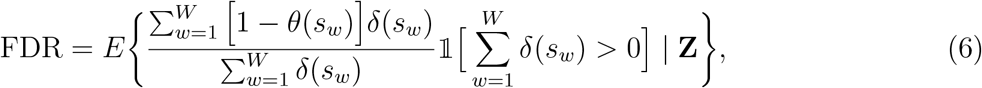

and the Bayesian false discovery exceedance^27^ (FDX) is the probability that the FDP exceeds some tolerance threshold *t*:

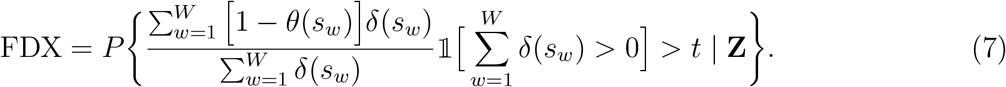

The FDX is useful in spatial contexts, where the FDR can be especially variable. Controlling the FDX instead of the FDR is akin to using a confidence interval instead of a point estimate; while it is a more conservative error quantity than FDR, its robustness to variable estimates can improve testing precision^41,42^.

Let *T*_*w*_ = *P* (*θ*(*s*_*w*_) = 0 |***Z***) be the probability that the null hypothesis is true at location *s*_*w*_ given ***Z***. Let *T*_(1)_ *≤ … ≤ T*_(*W*)_ be the ordered statistics *T*_*w*_ across all *W* windows, and *s*_(1)_, *…, s*_(*W*)_ be the corresponding windows. Then, we have

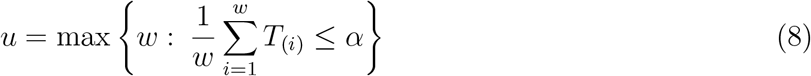

such that rejecting 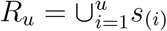 controls the FDR at level *α*. Likewise one controls the FDX by rejecting *R*_*u*_ when

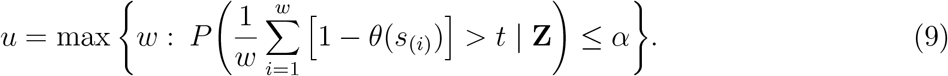

Because FDX involves 2 user-specified quantities–*t* and *α*–one typically says that the quantity in equation (9) achieves (*t, α*) *−*control. In all of our analyses, we recommend setting *t* equal to *α*, so for brevity refer to only a single parameter when discussing FDX control.

Since the hypothesis tests above are correlated across *w*, we adopt the statistical framework in Sun et al.^27^ for controlling false discoveries for spatially correlated tests. In brief, one first fits an appropriate spatial model (equation (4) using MCMC (Supplementary Section S1.3). The quantities in equations (8) and (9) are easily estimated from posterior MCMC samples (see Supplementary Section S1.4), and controlling the Bayesian quantities results in control of the frequentist analogs to FDR and FDX. We show how to adapt these quantities to better suit our setting in the subsequent section. Ultimately, this approach allows our NNGP model to produce rigorous false discovery control for differentially-interacting genomic regions while also accounting for complex spatial dependence in Hi-C data.

#### Integrating multiple scanning window sizes

One challenge in identifying differential regions from Hi-C data is that these regions are likely to come in various sizes for a given dataset, implying that there is not a one-size-fits-all scanning window size. We therefore fit multiple NNGP models, each with a transformed SCC sliding window statistic computed for a different window size as input. This strategy enables pinpointing differentially interacting regions of different sizes. Still, this approach introduces a new challenge for hypothesis testing, as multiple tests will be performed not only within the same study, but also several times at the same genomic location. That is, for example, one may fit 10 simultaneous NNGP models for 10 different sliding window sizes. Then, a given genomic region may be examined at least 10 times – once in each NNGP model.

To address this issue, we develop a procedure that incorporates information about the window size used in each analysis into the test for differential regions of Hi-C matrices. Our method extends the weighted false discovery rate (wFDR)^45^ and weighted false discovery exceedance (wFDX)^41^ into the Bayesian setting, where weights are naturally derived from the area of the scanning windows. We briefly describe the weighted testing procedures in our context. Suppose *J* different window sizes are considered, and let *W*_*j*_ be the number of scanning windows required to traverse a Hi-C matrix for window size *j*. Then, 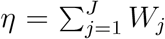 is the total number of windows tested across all analyses. For weights *a*_*i*_, *i ∈ {*1, *… η}*,

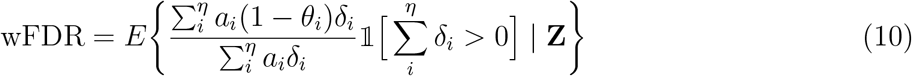

where *δ*_*i*_ is the *i*^*th*^ decision and *θ*_*i*_ is the truth at *δ*_*i*_’s location. Likewise, we introduce a Bayesian weighted FDX (wFDX) by extending the work of Perone Pacifico et al.^41^:

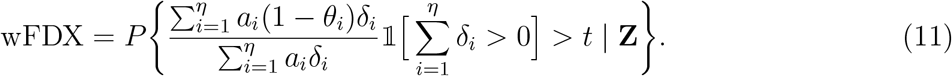

We describe how to calculate this quantity in Supplementary Section S1.4. These weighted quantities are designed such that one is penalized more for falsely rejecting a highly-weighted hypothesis. By assigning different weights to hypotheses that correspond to different window sizes, we can guard against falsely rejecting large regions. We weight each hypothesis by the area of the region the test corresponds to; that is, the weight for a hypothesis concerning a scanning window of size *B*_*w*_ is 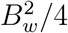.

This weighting is in fact a principled choice: Note that while our weighted hypothesis tests occur on a 1D NNGP, the rejections on this 1D process correspond to differential regions on a 2D Hi-C contact matrix. That is, we are ultimately interested in which entries of the Hi-C matrix will be marked as non-differential or differential. It turns out that weighting hypothesis tests based on the area of the sliding windows also controls FDR and FDX in 2D on the Hi-C contact map, stated in the following proposition (see *Methods* for definition of error rates, and Supplementary Section S2 for proof).

**Proposition 1**. *For any procedure that controls FDR along the 1D Gaussian process at level α, it also controls FDR in 2D on the Hi-C contact map*.

That controlling the wFDR (wFDX) in 1D implies FDR (FDX) control in 2D is directly corollary to this proposition. Therefore, though wFDR (wFDX) may be less easily interpreted relative to FDR (FDX), controlling these weighted quantities yields control of the usual FDR (FDX) in the 2D space, which is ultimately the space of interest.

### Simulations

To evaluate the performance of locdiffr against existing techniques, we used simulated polymer data based on the loop extrusion model to generate simulated Hi-C matrices^46,47^ (see *Methods*). Computationally generated Hi-C data from these polymer simulations provide a unique means to test methods designed for Hi-C data analysis, as these simulated data resemble real Hi-C data, yet are unmarred by experimental biases, and their ground truth is known.

We compared locdiffr, testing with both wFDR and wFDX, to existing tools multiHiCcompare^48^, selfish^49^, TopDom^17^, and edgeR^50^. MultiHiCcompare and selfish were designed specifically for differential analysis of Hi-C data, while TopDom calls TADs in a single experiment, and edgeR is typically used for differential gene expression analysis. We evaluated all methods based on the overlap of identified differential regions with the known true differences in the simulated data. (Implementation details for compared methods are in *Compared methods*).

We computed precision and recall pointwise on the Hi-C contact matrix (see *Methods*). The details of the simulation setup are shown in Table 1, and performance of each method under these simulation settings are in Fig. 2. Visualization of the rejected regions according to each method are in Supplementary Figs. S2-S10. Importantly, simulations 1 and 2 concern experimental conditions where some TAD boundaries are shared between conditions, while others are differential between conditions. In simulation 3, all TAD boundaries are shared between the two conditions, but there is a loss of intra-TAD interaction frequencies in one of the conditions compared to the other due to a simulated loss of insulation in one condition. For each simulation, we considered three sets of settings for other polymer model parameters, such as the direction of loop extrusion or TAD boundary strength (see *Polymer simulation overview*). Sample visualizations of each condition from each simulation are in supplementary Fig. S1.

**Table 1:** High-level simulation design for analysis of simulated Hi-C data. 1500 *×* 1500 Hi-C contact matrices were generated with 4 different sets of known TAD boundary locations (B1–B4). Across replicates within a given condition, all boundaries are the same (e.g., in Simulation 1 with contact radius 6, there are 7 replicates with boundary set B1, and 8 replicates with boundary set B2). A larger contact radius manifests as increase noise around the domain boundaries. Boundary sets B1 and B2 both have 51 boundary locations, of which 40 are shared between B1 and B2. Boundary set B3 has 72 boundaries, and boundary set B4 has 74 boundaries. A total of 46 boundary locations are shared between B3 and B4. In simulation 3, boundaries are all shared between the two conditions, but there is simulation-wide drop in intra-TAD interaction frequencies in condition 2 due to a simulated depletion of the TAD-insulating protein CTCF.

| Simulation | 1 |  |  | 2 |  |  | 3 |  |  |
| --- | --- | --- | --- | --- | --- | --- | --- | --- | --- |
| Boundary sets | B1/B2 | B1/B2 | B1/B2 | B3/B4 | B3/B4 | B3/B4 | B3/B3 | B3/B3 | B3/B3 |
| Contact radius | 6 | 8 | 10 | 6 | 8 | 10 | 6 | 8 | 10 |
| #reps., condition 1 | 7 | 7 | 7 | 7 | 7 | 7 | 7 | 7 | 7 |
| #reps., condition 2 | 8 | 8 | 8 | 4 | 4 | 4 | 9 | 9 | 9 |

**Figure 2:**
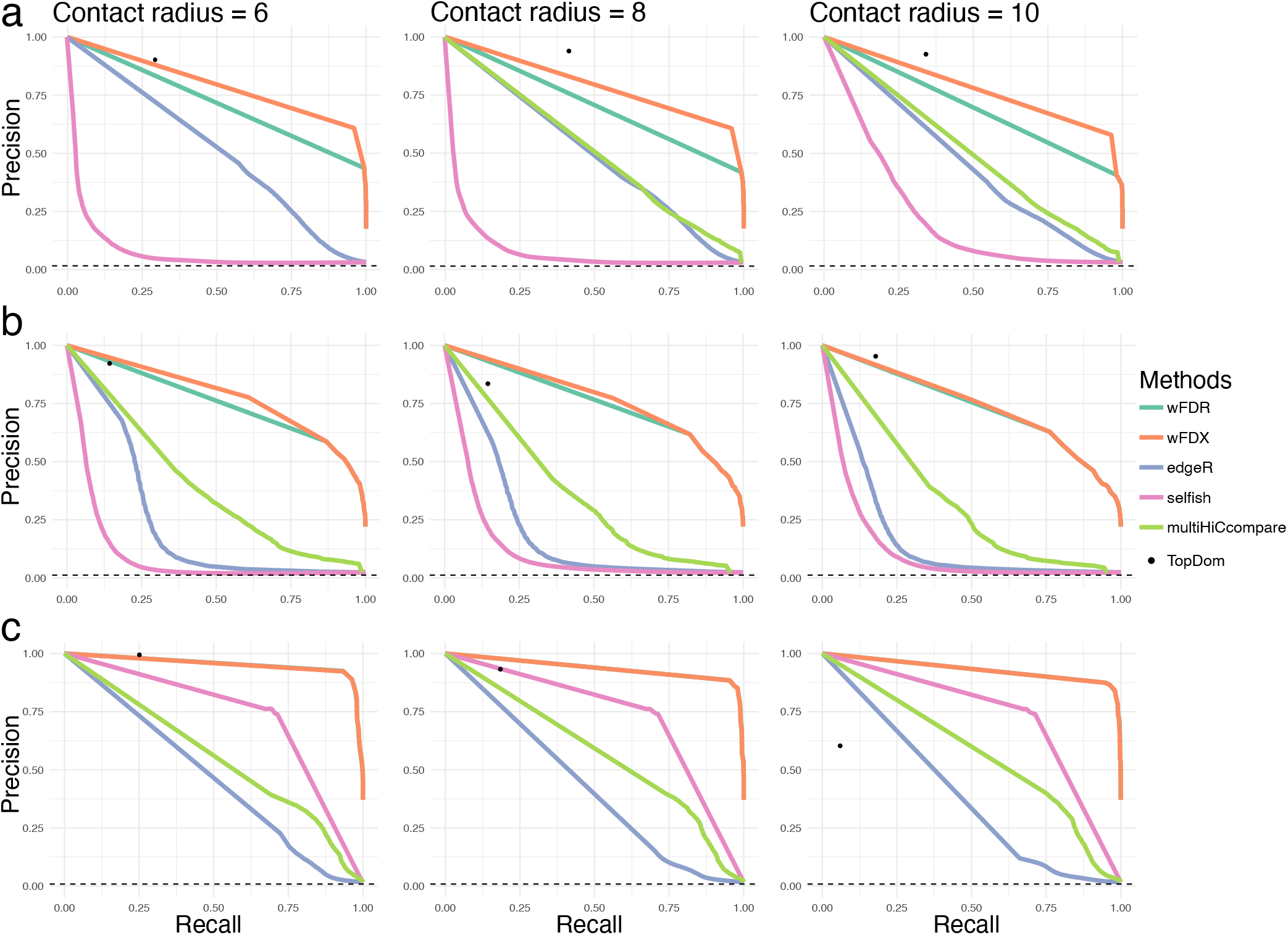
Precision-recall curves for all methods in **a**, simulation 1, **b**, simulation 2, and **c**, simulation 3. Each column corresponds to data simulated with different contact radii, where a larger contact radius manifests as more noise at simulated domain boundaries. The dashed black line represents the true proportion of pointwise differences between the two simulated conditions. The precision-recall curve for wFDR is sometimes obscured by that of wFDX, indicating similar rejections from both approaches. MultiHiCcompare failed to run under simulation 1 with contact radius 6.

Our results (Fig. 2) show that in general, locdiffr performed the best, especially when conducting tests with wFDX. As expected, multiHiCcompare – which adapts edgeR to address the 1D genomic distance dependence – always outperforms edgeR, affirming that treating all interactions as independent is inappropriate for Hi-C data. TopDom is frequently competitive, but its performance appears rather variable. Moreover, results from TopDom cannot be calibrated using *P*-values as all other compared methods can, yielding a single point on the precision-recall curve. Selfish is uniformly outperformed by all other methods in simulations 1 and 2, where domain boundaries vary between conditions. In simulation 3, where domain boundaries are shared, selfish outperforms both multiHiCcompare and edgeR.

### CTCF-depleted versus wild type in murine embryonic stem cells

CCCTC-binding factor (CTCF) and the cohesin complex are major actors in the formation of TADs and insulation of their boundaries^9,10,46^. Loss of CTCF does not eliminate TAD structure in mammalian cells, but it can cause a simultaneous increase in inter-TAD interactions and decrease in intra-TAD interactions^11,51^. Nora et al.^52^ explored how the degradation of CTCF affects chromatin structure in murine embryonic stem cells (mESCs) by depleting CTCF using the auxin-inducible degron (AID) system. We investigated the ability of locdiffr to find differences between CTCF-depleted (CTCF-AID) and wild type (WT) mESCs. We applied locdiffr to these Hi-C data (2 replicates from each condition) at 20 kb resolution. Because simulations showed superior performance with wFDX, we tested for significant differentially interacting genomic regions and controlled the wFDX at level .01.

Since differences in genome structure are expected to occur locally around eliminated CTCF binding sites, we expect locdiffr’s differential regions to coincide with CTCF binding sites that were sufficiently degraded in the CTCF-AID group. Figure 3a shows the average log_2_ ratio of CTCF ChIP-seq signals between the differential and non-differential regions that are identified with locdiffr (example track shot in Fig. S11). The identified differential regions show a significant enrichment for a loss of CTCF in the CTCF-AID cells. This result suggests that our approach is identifying regions exhibiting true biological differences.

**Figure 3:**
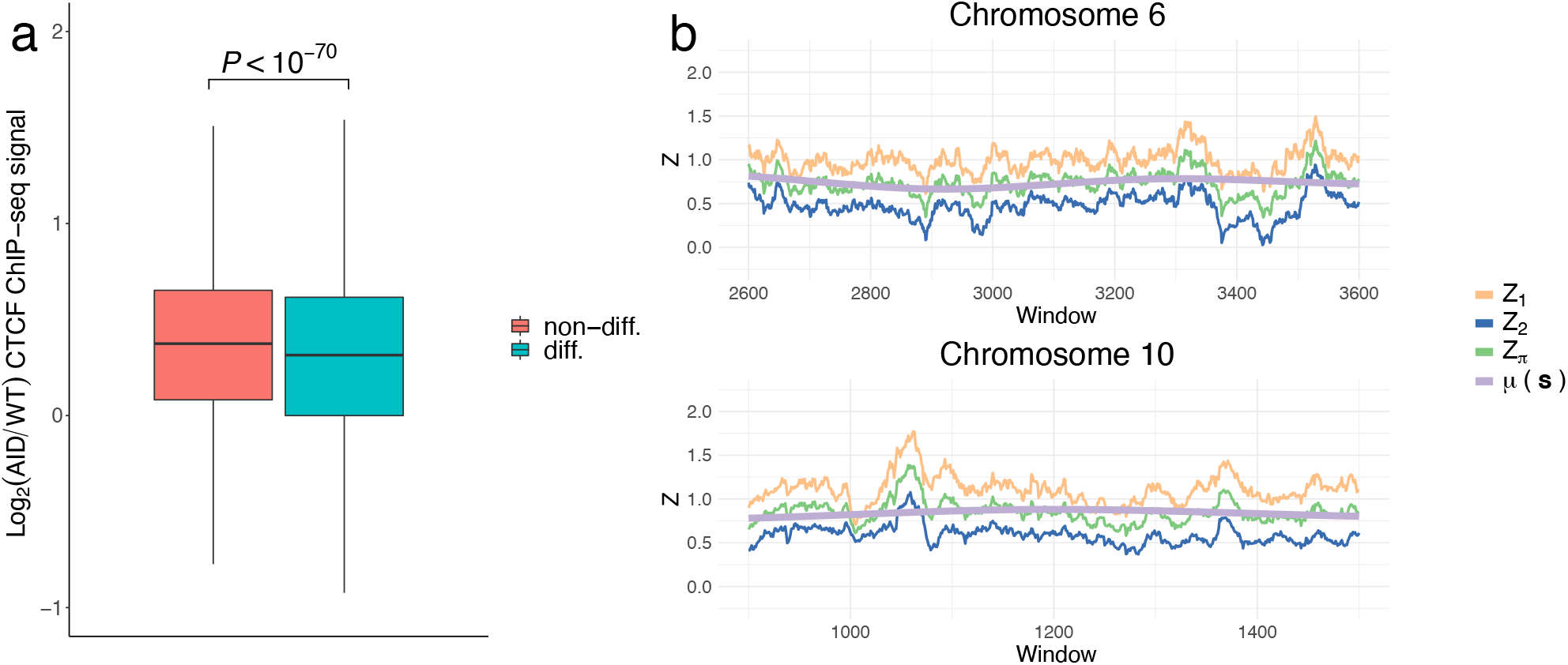
Differential analysis of Hi-C from wild type (WT) and CTCF depleted (AID) murine embryonic stem cells. **a**, 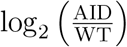 CTCF ChIP-seq signals in the significantly differential (i.e., rejected) and non-differential regions on the autosomes as described by locdiffr, divided into constituent 5 kilobase regions. There were 210,831 non-differential and 41,049 differential regions. The *P*-value was computed using a two-tailed Wilcoxon rank-sum test. Outliers were omitted from figure for ease of visualization. **b**, *z*-transformed SCC sliding window statistics (*Z*_1_ and *Z*_2_ for replicates 1 and 2, respectively) obtained with *B*_*w*_ = 40 from 2 example chromosomes plotted alongside the estimated mean function (*µ*(***s***)) and the average of all posterior draws (*Z*_*π*_, made with equation (19)) used to test which regions are significantly differential. *Z*_2_ was computed using the more lowly-sequenced experimental replicates.

Notably, the Hi-C data analyzed here have quite variable sequencing depth. For instance, for chromosome 10, there were *∼*8 million reads in both WT replicates, but *∼*15 million reads in one CTCF-AID replicate but only *∼*2 million reads in the other. As a consequence, Nora et al.^52^ pooled the Hi-C contact matrices across replicates prior to analysis, citing an inability to reproducibly call peaks in the data^53^. In contrast, with our approach, the lower sequencing depth for experimental replicates is reflected in our transformed SCC sliding window statistics (Fig. 3b, *Z*_2_ versus *Z*_1_). Despite this, the statistic computed using the lowly-sequenced replicate appears quite similar to the statistic computed from the more deeply sequenced replicates–i.e., the resulting processes generally possess the same peaks and valleys. This suggests that by borrowing strength from nearby bins using the sliding window statistic, our approach achieves some degree of robustness to a more lowly-sequenced experiment.

### Chd4-knockout versus wild type in murine cerebellar granule cells

Chromodomain Helicase DNA-binding protein 4 (Chd4) is a chromatin remodeling enzyme that is part of the Nucleosome Remodeling and Deacetylation (NuRD) complex. The NuRD complex is a transcriptional co-repressor that is known to influence chromatin accessibility and enhancer activity^54^, and it has frequently been tied to oncogenesis^55,56,57^. Recently, Goodman et al.^28^ studied WT and Chd4-knockout (KO) murine cerebellar cells, establishing a relationship between Chd4 and genome architecture in the mammalian brain. They identified that chromatin reorganization was tied to an increase in chromatin accessibility in the Chd4-KO cells as measured by DNaseI-seq genome-wide. We used locdiffr to analyze all autosomes of these Hi-C data (3 replicates from each condition) at 40 kb resolution, testing for differential genomic regions with control of the wFDX at level .01. By integrating with DNaseI- and ChIP-seq data (normalized to reads per million), we illustrate that our identification of significantly differentially interacting genomic regions complements these original findings.

#### Knockout of Chd4 results in genome-wide changes in epigenetic modifications

We first examined how DNaseI-seq and H3K4me1 and H3K27ac ChIP-seq signals changed between WT and KO groups, then compared these changes in the differential and non-differential regions. Results show strong evidence for an increase in chromatin accessibility and histone modification H3K4me1 within the regions with significant differences in their interaction frequencies genome-wide. In agreement with previous findings however^58^, large changes in H3K27ac are not evident using a broad, genome-wide approach (Fig. 4a). This pattern of epigenetic marks may be consistent with a transition of genomic regions, respectively, from inactive or primed enhancer states to primed or active enhancer states^59,60^. This comports with previous work that showed Chd4 in murine cerebellar cells functions as a promoter decommissioner, the loss of which may give rise to enhancer activation^58^. These results suggest that locdiffr identifies differential regions from Hi-C data that relate to biological function.

**Figure 4:**
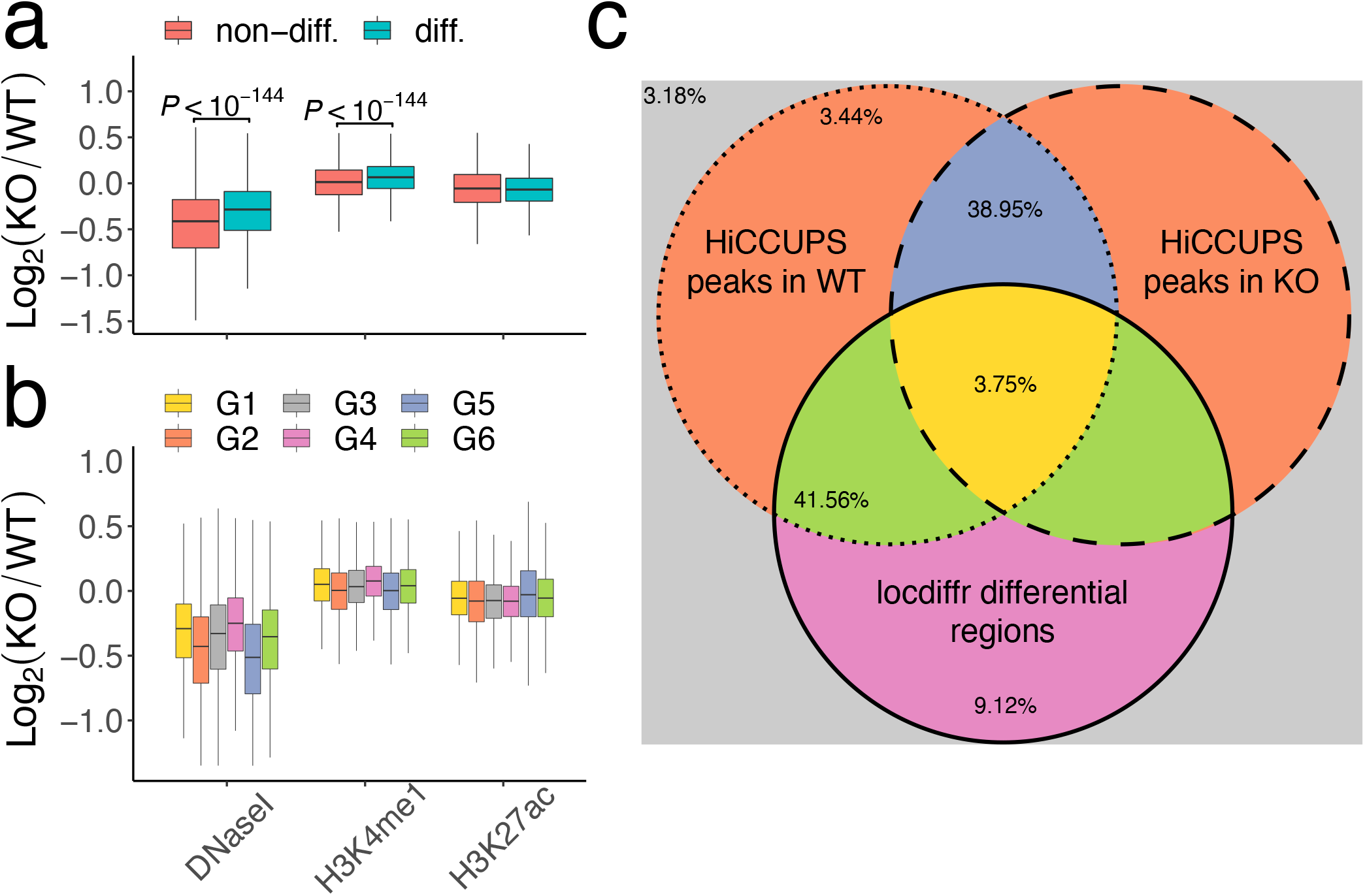
Comparison of enrichment of DNase- and ChIP-seq signals in wild type (WT) and knockout (KO) cells across the autosomes. **a**, 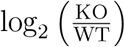 DNaseI-seq, and H3K4me1 and H3K27ac ChIP-seq signals, stratified into rejected and null regions as described by locdiffr and **b**, stratified into groups by comparing locdiffr’s differential and non-differential regions to those regions identified with HiCCUPS, divided into constituent 5 kilobase regions. Locdiffr produced 99,258 non-differential and 180,909 differential regions. *P*-values were computed using a two-tailed Wilcoxon rank-sum test. Outliers were omitted from figure for ease of visualization. Signals taken only from within peak regions produce boxplots showing similar trends as **a**–**b** (supplementary Fig. S12). **c**, Illustration of partitioning of genomic regions into *G*1 *− G*6. Differential regions according to our approach are marked by the solid circle. Regions proximal to a loop anchor identified with HiCCUPS in Goodman et al. in wild type (WT) and Chd4-knockout (KO) cells are marked by the dotted and dashed circles, respectively. Some genomic regions are not proximal to a loop anchor in either condition. Colors are matched to the legend in **b**. Annotated percentages indicate the percentage of the autosomal regions assigned to each group.

#### Analysis of differential regions complements the analysis of differential peaks

Next, we examined how our findings compared and contrasted against some of the analyses of the Hi-C data originally done by Goodman et al.^28^. Briefly, in their work, they used HiCCUPS^61^ to identify loop anchors in each condition, then described these anchors as either overlapping, or specific to one of the WT or KO conditions. To make a comparison, we partitioned genomic loci into several groups, based on both the significant local differences in chromatin architecture called by locdiffr and the HiCCUPS loop anchor status.

The partition is as follows: Let *N*_*L*_ and *D*_*L*_ be the sets of loci in the non-differential and differential regions, respectively, as identified by locdiffr. Then, let *N*_*H*_ and *D*_*H*_ be regions marked by loop anchors that respectively overlap in both conditions, or are specific to just one condition, as identified by Goodman et al. using HiCCUPS. Note that while the set *N*_*L*_ *∪ D*_*L*_ covers all modeled loci, the set *N*_*H*_ *∪ D*_*H*_ only covers regions associated with loop anchors in at least one condition. Because of this, to portray a complete picture of the results, we examined epigenetic signals by partitioning the genome into 6 different regions. These sets allow us to compare how the differential regions identified with locdiffr and HiCCUPS correlate with epigenetic changes (Fig. 4b), while also stratifying by the presence of loop anchors. The sets are:

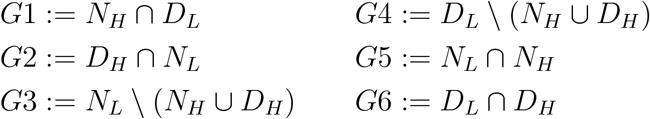

Here, *A \ B* is the set difference of *A* and *B*. The majority (80.51%) of autosomal regions fall into the groups where locdiffr and HiCCUPS agree (*G*5 and *G*6). An illustration of all groups, and their relative sizes, are in Fig. 4c. A sample track shot displaying the locations of each group in the partition is in Fig. 5.

**Figure 5:**
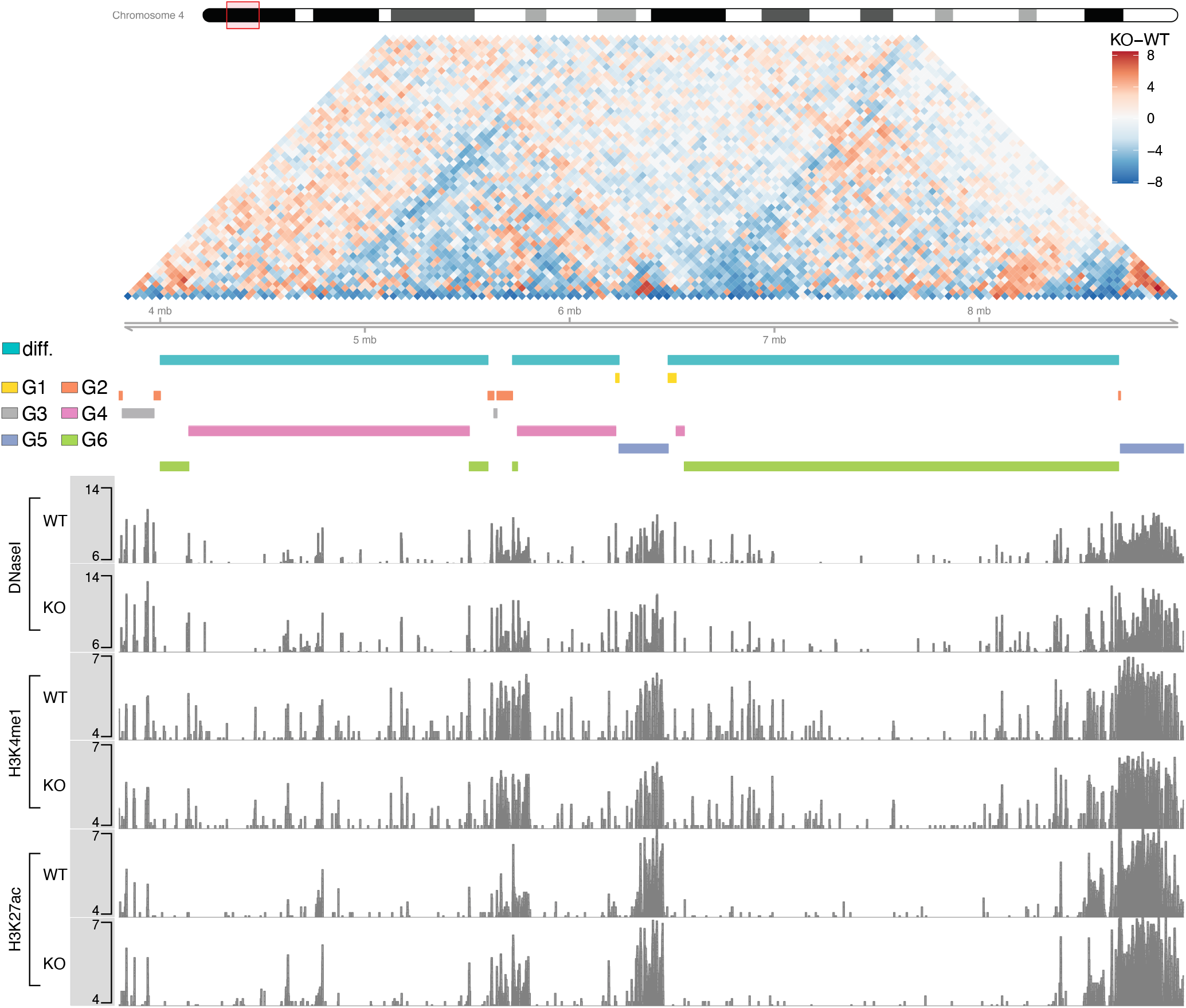
Sample track shot of significantly differential regions on chromosome 4 in the Chd4-knockout analysis, shown alongside analyzed Hi-C data. Track signals (for ChIP-seq) and differences in interactions between wild type and Chd4-knockout experiments (for Hi-C) are on the log_2_ scale. Regions are annotated by their partition, colored to match Fig. 4c. Teal bars annotate all locdiffr differential regions.

Again, most striking in Fig. 4b are the differences evident among differential enrichment in DNaseI and H3K4me1 signals. In the case of DNaseI, the regions identified as differential by both approaches in *G*6 show an increase in chromatin accessibility in the KO cells when compared against group *G*5, the set of regions both methods called non-differential. Contrasting *G*5 and *G*6, the loci in group *G*1 exhibit some disagreement between locdiffr and HiCCUPS; these loci were called as different in architecture by locdiffr, whereas the loop anchors were called as the same by HiCCUPS. Remarkably, the *G*1 loci show an even greater increase in accessibility than the *G*6 loci, suggesting that locdiffr identified truly differential domains that HiCCUPS labeled as non-differential. Meanwhile, the smallest increase in chromatin accessibility over group *G*5 was found in group *G*2, suggesting that the set of differential peaks called with HiCCUPS, which locdiffr called non-differential, is populated by many regions not exhibiting changes in chromatin accessibility. Further, HiCCUPS-specific differential regions (*G*2) do not show enrichment in differential H3K4me1 over baseline group *G*5. However, a gain in differential H3K4me1 is evident in regions locdiffr identified as differential (*G*1, *G*4, and *G*6).

Quite interestingly, we also noticed an increase in H3K27ac in group *G*5 over all other groups. As exemplified in Fig. 5, we can observe increases in H3K27ac signal abutting the boundaries of locdiffr’s differential regions. This is in contrast to H3K4me1, which tends to instead exhibit increased signal in the KO population within the interior of the differential regions (*G*1, *G*4, and *G*6).

Even though no difference was detected in the distributions of H3K27ac signals between the differential and non-differential regions called by locdiffr (Fig. 4a), when only the loop anchors (from HiCCUPS) are examined in the nondifferential regions (*G*5), they show significantly higher levels of this histone modification in the KO group (Fig. 4b). Given that knockout of Chd4 is associated with increased enhancer activity^28^, our results suggest that the loop anchors with nearly unchanged chromatin interaction after Chd4 knockout (i.e. those in common by HiCCUPS and confirmed as non-differential by locdiffr) may still be engaged in enhancement as indicated by their retention of significantly higher H3K27 acetylation.

Altogether these results show that many biologically functional differential interaction regions may not necessarily be associated with loop anchors, suggesting that it is insufficient to only search for peaks in Hi-C data. Instead, locdiffr effectively identifies these differential interaction regions, complementing peak-calling data. Integrating both methods can provide a richer understanding of the relationship between 3D genome architecture and genome-wide epigenetic signals.

## Discussion

We introduced a statistical methodology, called locdiffr, to detect local changes in genome architecture between Hi-C samples. We contend that locdiffr is more appropriate for controlling false discoveries when testing for significant differences, as it more directly accounts for complex dependence present in Hi-C data. We validated this claim using polymer model-based simulations, finding that locdiffr outperforms existing methods in both power and precision, especially when choosing to control the wFDX. We also demonstrated locdiffr’s approach on two WT versus treatment datasets, showing that differential regions identified by our approach correlate with epigenetic modifications, and that our findings differ from and complement the findings of peak-calling tool HiCCUPS^61^. We further illustrated locdiffr’s robustness in the common setting when experimental replicates are variably sequenced.

Like some peak-based methods (e.g., multiHiCcompare), but unlike TAD callers, locdiffr uses experimental replicates to incorporate information about variance into the modeling and testing procedure, rather than simply pooling the data across replicates. The leveraging of between-replicate variance may be extended to within-replicate variance in future enhancements of our approach. Specifically, adapting locdiffr to also identify irreproducible regions by examining concordance among within-condition samples, such that irreproducible sites are excluded from hypothesis testing, could boost our method’s precision.

## Methods

### Polymer simulation overview

An overview of our simulation strategy is presented here (details in supplementary section S3). Polymer simulations were performed using the polychrom library^62^, which is a wrapper around OpenMM^63^. OpenMM is an R package for performing efficient GPU-assisted molecular dynamics or Langevin dynamics simulations. We simulated 30 Megabases (Mb) of chromatin fiber as a sequence of 15,000 monomers, each representing 2 kb of DNA. Each monomer was a sphere of 20 nanometers (nm) in diameter. For each arrangement of TADs we simulated at least 12 independent simulations, each containing 4 copies of the system. Each simulation included 100,000 steps of loop extrusion, which translates to 60,000,000 steps of molecular dynamics. Conformations were recorded every 20 steps of molecular dynamics.

Different replicates were simulated by randomly assigning a number between 1 and 10 to each simulation, and pooling simulations with different numbers. This created replicates with different coverage and different amount of noise, consistent with how replicates appear in Hi-C experiments. Contact maps were calculated using polychrom at 3 different contact radii: 6, 8, and 10 monomers (120nm, 160nm and 200nm). All pairs of monomers that are closer than the contact radius are counted as contacts. For display and comparison with Hi-C, contact maps were coarse-grained to 20 kb resolution by summing all the pixels in a 10 *×* 10 pixel (20 *×* 20 kb) squares.

#### Arrangements of simulated TADs

To compare the examined methods, we generated several artificial arrangements of TADs. For the first simulation, we created a system made of 62 TADs with a mean TAD size of 250 monomers. TAD sizes were selected from an exponential distribution with rate *λ* = 150, and only TAD sizes more than 100 were picked. TAD sizes were rounded to the nearest value of 10, since the heatmap would be later coarsegrained by a factor of 10. The resulting process created TADs with an average size of about 250 monomers, ranging from 110 monomers to 620 monomers. To simulate two conditions with different TAD arrangements, we selected 51 TAD boundary at random for each of the two conditions. As a result, the two conditions were different at 22 boundaries.

For the second simulation, we simulated a system of 60 TADs, arranged as 6 groups of 10 TADs. The length of the TADs were selected from a uniform distribution between 190 and 330 monomers, again only taking multiples of 10. From this arrangement of TADs, we simulated two conditions with different changes between the two conditions. Changes were mild in the first group of 10 TADs, and become progressively stronger in later groups. Within each group of 10 TADs, the following modifications were made: The first TAD in each group had a bidirectional boundary added in the center, but only in one randomly chosen condition. The third TAD had a uni-directional boundary added in the center in a randomly chosen condition. The fifth TAD had two uni-directional boundaries added at 1*/*3 and 2*/*3 of the TAD in a randomly chosen condition; the first boundary pointed right (at the second boundary), and the second boundary pointed left (at the first boundary). The seventh TAD in each group had one of its two boundaries shifted in a randomly chosen condition. The ninth TAD had both boundaries shifted in the same direction in a randomly chosen condition.

The added boundaries had different *strengths*, defined as the probability that the boundary stops a loop extruding factor (LEF). The strengths of the added boundaries were 0.1, 0.2, 0.3, 0.5, 0.7, 0.9 in groups 1 through 6, while the strengths of normal boundaries were 1. Shifts of boundaries for TADs 7 and 9 were 10, 20, 30, 40, 50, 60 monomers in groups 1 through 6.

For the third simulation, we took the first condition used in the second simulation, and kept the TAD arrangements the same. However, we reduced the number of LEFs by a factor of 5, thus making loop extrusion weaker.

### Implementation details

#### locdiffr

Window sizes, used to compute the SCC scan statistics, were always selected evenly along a grid (Table 2). In simulations, since true differential region sizes are known, we selected the window sizes to span a grid from the smallest known differential region to the largest known differential region.

**Table 2:** MCMC details for all locdiffr analyses.

| Data | min. window | max. window | # windows | burn-in | total iterations |
| --- | --- | --- | --- | --- | --- |
| Simulation 1 | 28 | 139 | 20 | 10,000 | 25,000 |
| Simulation 2 | 13 | 84 | 20 | 10,000 | 25,000 |
| Simulation 3 | 6 | 35 | 20 | 10,000 | 25,000 |
| CTCF-AID data | 20 | 100 | 5 | 5,000 | 15,000 |
| Chd4-KO data | 10 | 200 | 20 | 5,000 | 15,000 |

Locdiffr requires pairing samples from each condition. Yet, in the simulated data, there are unequal numbers of replicates from each condition. Therefore, our final sample size was equal to the number of replicates from the condition with fewer replicates. Samples are paired based on sequencing depth, e.g., the sample with the most counts in the first condition is paired with the sample with the most counts from the second condition. The SCC score can be sensitive to sequencing depth, so to avoid this source of uncertainty, downsampling is done within each pair such that the number of reads for each sample within a pair is matched. In cases where there are more replicates from one condition than another, to omit extra replicates, we removed those with the lowest number of counts. For all other compared methods, no replicates were omitted.

A brief summary of MCMC iterations used to fit the NNGP are in Table 2. In both the simulations as well as each chromosome analyzed from the 2 real datasets, we used 1 nearest neighbor for each bin and set *p* = *{W/*300, 4*}* for cubic B-splines. In both simulations, we used 10,000 posterior samples in the set *l ∈ {*5, 002, 5, 004, *…*, 25, 000*}* to estimate wFDR and wFDX. For the analysis of each chromosome of the CTCF-AID and Chd4-KO data, we used 10,000 posterior samples in the set *l ∈ {*5, 001, 5, 002, *…*, 15, 000*}*. To compute wFDX, we always set *t* = *α* and used 100 bootstrap replicates.

#### Compared methods

##### multiHiCcompare

We implemented multiHiCcompare using the software package as advised in documentation. We created a Hi-C experiment with the function `make_hicexp` at default settings, and pre-processed the data with the cyclic loess normalized technique. We fit the generalized linear model with the `hic_glm` function, and tested for significantly different interactions using the quasi-likelihood *F*-test and BH-adjusted *P*-values.

##### Selfish

Selfish does not request that the data be normalized. Therefore, we simply ran Selfish at default settings, and evaluated significance based on BH-adjusted *P*-values.

##### TopDom

Since TopDom does not use replicates, we first summed over all simulated data within each simulated condition. We called TADs in each condition with TopDom using a window size of 10. TopDom returns the start and stop locations of all called TADs. We called a region differential between conditions 1 and 2 if the region fell within a TAD in one condition, but not in the other.

##### edgeR

Unlike other compared methods, edgeR was developed for RNA-seq analyses. It is a key component in multiHiCCompare for identifying differential Hi-C bins. It treats all bins of the Hi-C contact matrix as independent, regardless of spatial proximity. To apply this method, we first transformed the contact matrix to a vector, and downsampled them to equal read counts. We ran edgeR using the function `glmQLFit`, and identified significantly differential interactions using the quasi-likelihood *F*-test and BH-adjusted *P*-values.

#### Evaluating error rates

We evaluated error rates pointwise between estimated and true differences between simulated Hi-C contact matrices from two conditions. For a given simulation, let *S*_*P*_ be the set of entries in an upper-triangular matrix of dimension *W × W*, excluding the diagonal, which corresponds to true pointwise differences between two simulated condition. Let *ŜP* similarly define the pointwise differences estimated by a given method. Then, for all methods, we define

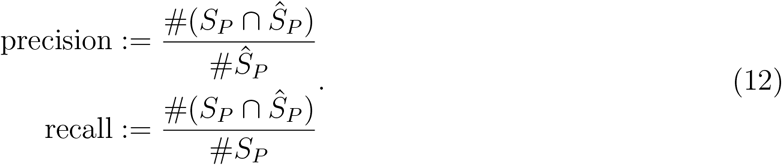

#### Enrichment analyses

To perform enrichment analyses, we first classified al l bins as non-differential or differential with locdiffr. Because loci annotated by both locdiffr and HiCCUPS are of various sizes, we first sought to create more regularly sized partitions of the loci. Therefore, we partitioned each bin into 5 kb sub-bins. For the Chd4-KO data, we constructed groups *G*1 *− G*6 by comparing these sub-bins with those of Goodman et al.^28^ using BEDtools v2.27.1^64^. To make Fig. 3a and Fig. 4a–b, we computed the average signal within each resulting region of length greater than or equal to 0.25 kb using `bigWigAverageOverBed`^65^. We similarly used `bigWigAverageOverBed` to compute the average log_2_ fold change in supplementary Figs. S12 and S13. A common custom blacklist region, including heterochromatic regions, was exluded from all enrichment analyses. Track shots were made in R v3.6.2^66^ using the package Gviz v1.30.3^67^.

## Availability

Locdiffr is implemented in an R package, freely available on GitHub under an Artistic-2.0 license (https://github.com/hillarykoch/locdiffr). Examined datasets are available at NCBI’s Gene Expression Omnibus (https://www.ncbi.nlm.nih.gov/geo/)^68^. CTCF-AID data are available under accession code GSE98671. Chd4-KO data are available under accession code GSE138822.

## Acknowledgments

The authors are grateful for the support from their funding agencies: NHGRI pre-doctoral fellowship 1F31HG010574 and NIH training grant T32 GM102057 (CBIOS training program to The Pennsylvania State University) to H.K.; support from the NIH Common Fund 4D Nucleome Program (DK107980) to M.I.; NIDDK grant R24DK106766 to R.C.H.; and NIGMS grant R01GM109453 to Q.L..

