## Supplementary Materials for "Detecting local changes in chromatin architecture with false discovery control"

### Supplementary Material

#### S1 MCMC details

##### S1.1 Bounding the spatial range

The number of Hi-C experiments available for comparative analysis is likely to be small ($n < 5$), and can therefore make parameter inference quite challenging. This is especially true for the spatial range $r$, a measure of the strength of spatial lag, which can be difficult to estimate even with large sample sizes. By predefining suitable bounds on $r$, parameter inference may become simpler. A natural lower bound on $r$ is 1, given the discrete nature of the sliding window and that $r = 0$ would imply no spatial dependence. We obtain an upper bound on $r$ by examining the decay of autocorrelation in the data. First, define $Z_j^\star(\mathbf{s})$ as the Gaussian process comprised of pointwise averages across $\mathbf{Z}_j$ at spatial locations $\mathbf{s}$ in analysis $j$, and $\bar{Z}_j^\star(\mathbf{s})$ as the average of $Z_j^\star(\mathbf{s})$, such that

$$\Gamma_\gamma(Z_j^\star(\mathbf{s})) = \frac{\sum_{w=1}^{W-\gamma}\left[Z_j^\star(s_w) - \bar{Z}_j^\star(\mathbf{s})\right]\left[Z_j^\star(s_{w+\gamma}) - \bar{Z}_j^\star(\mathbf{s})\right]}{\sum_{w=1}^{W}\left[Z_j^\star(s_w) - \bar{Z}_j^\star(\mathbf{s})\right]^2} \tag{13}$$

is the autocorrelation observed in process $Z_j^\star(\mathbf{s})$ at spatial lag $\gamma$. Then the function $\psi$ finds the effective lag

$$\psi\left[Z_j^\star(\mathbf{s})\right] = \min_{\gamma\in\{1,\ldots,W_j\}}\left\{\Gamma_\gamma(Z_j^\star(\mathbf{s})) < 0.05 \text{ and } \Gamma_\gamma(Z_j^\star(\mathbf{s})) < \Gamma_{\gamma+1}(Z_j^\star(\mathbf{s}))\right\}. \tag{14}$$

This function returns the first lag whose autocorrelation is less than 0.05, before the autocorrelation begins increasing at larger lags.

##### S1.2 Selecting the covariance function and the number of neighbors

In this work, we modeled the transformed sliding window statistic approximately as a Gaussian process with an exponential covariance function. In 1-dimensional discrete space, the exponential covariance function simplifies to that induced by a Gaussian AR(1) time series. Inference on such a process is implictly accomplished by fitting a 1-dimensional NNGP with $k = 1$ neighbors.

Partial autocorrelation functions of the transformed sliding window statistics from the simulation studies (Fig. S14) and both real data analyses (Fig. S15 and S16) support that the exponential covariance function is a suitable modeling choice, since the partial autocorrelation significantly decays after lag 1.

##### S1.3 MCMC sampling

We estimate the parameters in the NNGP model using a Metropolis-within-Gibbs sampling scheme. Gibbs updates are available for $\boldsymbol{\beta}$ and $\sigma^2$, while Metropolis steps must be executed to sample $r$ and $\epsilon^2$. An overview of the sampling procedure is in Algorithm 1. For simplicity, we omit subscript $j$ indicating a specific Markov chain for a given scanning window size.

Let $\tilde{C}(\mathbf{s}, \mathbf{s};\, r) = \tilde{C}$ be the exponential correlation function across spatial locations $\mathbf{s}$ parameterized by the spatial range $r$. Then,

**Algorithm 1** Sample NNGP parameters with $L$ MCMC iterations

Initialize parameters at $\boldsymbol{\beta}^{(0)}$, $\sigma^{2(0)}$, $r^{(0)}, \epsilon^{2(0)}$
Define tuning parameters $\tau_r$ and $\tau_\epsilon$
**for** $l$ in 1 to $L$ **do**
    Draw $\boldsymbol{\beta}^{(l)} \sim \pi\Big(\boldsymbol{\beta} \mid \sigma^{2(l-1)},\, r^{(l-1)},\, \epsilon^{2(l-1)}\Big)$
    Draw $1/\sigma^{2(l)} \sim \pi\Big(1/\sigma^2 \mid \boldsymbol{\beta}^{(l)},\, r^{(l-1)},\, \epsilon^{2(l-1)}\Big)$
    Propose $r^\star \sim \mathrm{N}(r^{(l-1)},\, \tau_r)$
    Compute ratio $R_r = h(r^\star)/h(r^{(l-1)}) = \exp\Big[\log h(r^\star) - \log h(r^{(l-1)})\Big]$
    **if** $R_r \geq 1$ **then**
        $r^{(l)} \leftarrow r^\star$
    **else**
        $r^{(l)} \leftarrow r^\star$ with probability $R_r$, otherwise $r^{(l)} \leftarrow r^{(l-1)}$
    Propose $1/\epsilon^{2\star} \sim \mathrm{N}(1/\epsilon^{2(l-1)},\, \tau_\epsilon)$
    Compute ratio $R_\epsilon = h(1/\epsilon^\star)/h(1/\epsilon^{(l-1)}) = \exp\Big[\log h(1/\epsilon^\star) - \log h(1/\epsilon^{(l-1)})\Big]$
    **if** $R_\epsilon \geq 1$ **then**
        $1/\epsilon^{(l)} \leftarrow 1/\epsilon^\star$
    **else**
        $1/\epsilon^{(l)} \leftarrow 1/\epsilon^\star$ with probability $R_\epsilon$, otherwise $1/\epsilon^{(l)} \leftarrow 1/\epsilon^{(l-1)}$

$$\begin{aligned}
&\pi\Big(\boldsymbol{\beta} \mid \sigma^{2(l-1)},\, r^{(l-1)},\, \epsilon^{2(l-1)}\Big) := \\
&\qquad \mathrm{N}\Bigg(\Big[\frac{n}{\sigma^{2(l-1)}} X^{\mathrm{T}}[\tilde{C}^{(l-1)} + \epsilon^{2(l-1)}\mathrm{I}]^{-1}X + \frac{1}{9}\mathrm{I}\Big]^{-1}\Big[\frac{n}{\sigma^{2(l-1)}} X^{\mathrm{T}}[\tilde{C}^{(l-1)} + \epsilon^{2(l-1)}\mathrm{I}]^{-1}Z^\star(\mathbf{s})\Big], \\
&\qquad\qquad \Big[\frac{n}{\sigma^2} X^{\mathrm{T}}[\tilde{C}^{(l-1)} + \epsilon^{2(l-1)}\mathrm{I}]^{-1}X + \frac{1}{9}\mathrm{I}\Big]^{-1}\Bigg)
\end{aligned} \tag{15}$$

and

$$\begin{aligned}
&\pi\Big(1/\sigma^2 \mid \boldsymbol{\beta}^{(l)},\, r^{(l-1)},\, \epsilon^{2(l-1)}\Big) := \\
&\qquad \mathrm{Gamma}\Bigg(\frac{nW}{2} + 2,\, \frac{\sum_{i=1}^{n}(Z_i(\mathbf{s}) - X\boldsymbol{\beta}^{(l)})^{\mathrm{T}}(\tilde{C}^{(l-1)} + \epsilon^{2(l-1)}\mathrm{I})^{-1}(Z_i(\mathbf{s}) - X\boldsymbol{\beta}^{(l)})}{2} + 2\Bigg).
\end{aligned} \tag{16}$$

The acceptance ratio $R_r$ is

$$
\begin{aligned}
R_r = \exp\Bigg\{ & \frac{W}{2}\Big[\log|\sigma^{2(l)}(\tilde{C}^{(l-1)} + \epsilon^{2(l-1)}\mathrm{I})| - \log|\sigma^{2(l)}(\tilde{C}^{\star} + \epsilon^{2(l-1)}\mathrm{I})|\Big] \\
& + \frac{1}{2\sigma^{2(l)}}\Big[\sum_{i=1}^{n}(Z_i(\mathbf{s}) - X\boldsymbol{\beta}^{(l)})^{\mathrm{T}}(\tilde{C}^{(l-1)} + \epsilon^{2(l-1)}\mathrm{I})^{-1}(Z_i(\mathbf{s}) - X\boldsymbol{\beta}^{(l)}) \\
& - \sum_{i=1}^{n}(Z_i(\mathbf{s}) - X\boldsymbol{\beta}^{(l)})^{\mathrm{T}}(\tilde{C}^{\star} + \epsilon^{2(l-1)}\mathrm{I})^{-1}(Z_i(\mathbf{s}) - X\boldsymbol{\beta}^{(l)})\Big]\Bigg\}
\end{aligned} \tag{17}
$$

and the acceptance ration $R_\epsilon$ is

$$
\begin{aligned}
R_\epsilon = \exp\Bigg\{ & \frac{W}{2}\Big[\log|\sigma^{2(l)}[\tilde{C}^{(l)} + \epsilon^{2(l-1)}\mathrm{I}]| - \log|\sigma^{2(l)}[\tilde{C}^{(l)} + \epsilon^{2\star}\mathrm{I}]|\Big] \\
& + \frac{1}{2\sigma^{2(l)}}\Big[\sum_{i=1}^{n}(Z_i(\mathbf{s}) - X\boldsymbol{\beta}^{(l)})^{\mathrm{T}}(\tilde{C}^{(l)} + \epsilon^{2(l-1)}\mathrm{I})^{-1}(Z_i(\mathbf{s}) - X\boldsymbol{\beta}^{(l)}) \\
& - \sum_{i=1}^{n}(Z_i(\mathbf{s}) - X\boldsymbol{\beta}^{(l)})^{\mathrm{T}}(\tilde{C}^{(l)} + \epsilon^{2\star}\mathrm{I})^{-1}(Z_i(\mathbf{s}) - X\boldsymbol{\beta}^{(l)})\Big] \\
& + \log\Big[g(1/\epsilon^{2\star})\Big] - \log\Big[g(1/\epsilon^{2(l-1)})\Big]\Bigg\}
\end{aligned} \tag{18}
$$

where $g(x)$ is the density of Gamma-distributed $x$ with shape and scale parameters both equal to 2.

#### S1.4 Estimating false discovery controlling quantities from posterior MCMC samples

It is straightforward to estimate the quantities in equations (10) and (11) using the posterior MCMC samples. First, define at iteration $l$ in analysis $j$

$$
Z_{\pi,j}^{(l)}(\mathbf{s}) \sim \mathrm{NNGP}\left(Z_j^{\star}(\mathbf{s}),\, \sigma_j^{2(l)}\tilde{C}_j^{(l)}\right) \tag{19}
$$

as a newly sampled NNGP at spatial locations $\mathbf{s}$, with parameters $\boldsymbol{\beta}_j^{(l)}$, $\sigma_j^{2(l)}$, and $r_j^{(l)}$. The mean of $Z_j^{\star}(\mathbf{s})$ is the conditional mean of the newly sampled data given the observed data and parameters at iteration $l$, and the covariance is the conditional variance of the newly sampled data given the parameters at iteration $l$, assuming independence between the data $\mathbf{Z}$ and $Z_{\pi,j}^{(l)}$. This covariance is meant to reflect the underlying biological variance, while the nugget is omitted as it is intended to capture iid technical variablility. One can sample such a NNGP using an adaptation of Algorithm 4 in Finley et al. (2019)[38].

Next, let $\theta^{(l)}(s_{w,j}) = \mathbb{1}[Z_{\pi,j}^{(l)}(s_{w,j}) < \mu^{(l)}(s_{w,j})]$ be the indicator that the sampled NNGP lies in the rejection region at location $s_w$ on MCMC iteration $l$ in analysis $j$. Building upon previous work[27], we use these indicators to estimate wFDR and wFDX.

We define $\hat{T}_{w,j}$, $w \in \{1, \ldots, W_j\}$, $j \in \{1, \ldots, J\}$, as

$$
\begin{aligned}
T_{w,j} &= P(\theta(s_{w,j}) = 0 \mid \mathbf{Z}_j) \\
&= \int_{Z_{\pi,j}} \int_{\mu_j} \mathbb{1}\Big[Z_{\pi,j}(s_{w,j}) < \mu_j(s_{w,j})\Big] \pi\Big[Z_{\pi,j}(\mathbf{s}),\, \boldsymbol{\beta}_j \mid \mathbf{Z}_j,\, \sigma_j^2,\, r_j\Big] d\mu_j dZ_{\pi,j}
\end{aligned} \tag{20}
$$

which is estimable by

$$
\hat{T}_{w,j} = \frac{1}{L} \sum_{l=1}^{L} [1 - \theta^{(l)}(s_{w,j})] \tag{21}
$$

where $\mathbf{Z}_j$ is the data from analysis $j$, and $s_{w,j}$ is the $w^{\text{th}}$ window in analysis $j$. Ordering $\hat{T}_{1,1}, \ldots, \hat{T}_{W_J,J}$ as $\hat{T}_{(1)}, \ldots, \hat{T}_{(\eta)}$, one can substitute these values into equation (10) to obtain wFDR control at level $\alpha$. One can similarly use these quantities to obtain $(t, \alpha)-$control with wFDX. Algorithms 2 and 3 explicitly describe procedures for controlling wFDR and wFDX. Note that for the bootstrapping in Algorithm 3, it is essential that calculations are based on the same set of windows in each replicate. That is, ordering of statistics $T_i^{boot}$ are done based on only 1 set of posterior draws $Z_{\pi,j}^{(l)}(\mathbf{s})$.

**Algorithm 2** Control wFDR at level $\alpha$ using $L$ posterior MCMC samples.

1: Compute $\hat{T}_{(1)}, \ldots, \hat{T}_{(\eta)}$ and corresponding weigths $a_{(1)}, \ldots, a_{(\eta)}$
2: **for** $u'$ in $1 \ldots \eta$ **do**
3:     $\text{wFDR}(u') \leftarrow \frac{\sum_{i=1}^{u'} a_{(i)} \hat{T}_{(i)}}{\sum_{i=1}^{u'} a_{(i)}}$
4: Compute $u = \text{argmax}_{u' \in \{1,\ldots,\eta\}} \Big\{ \text{wFDR}(u') \leq \alpha \Big\}$
5: Reject hypotheses at the windows corresponding to statistics $\hat{T}_{(1)}, \ldots, \hat{T}_{(u)}$

**Algorithm 3** Control wFDX at level $(t, \alpha)$ using $L$ posterior MCMC samples.

1: Compute $\hat{T}_{(1)}, \ldots, \hat{T}_{(\eta)}$ and corresponding weigths $a_{(1)}, \ldots, a_{(\eta)}$ and assume for ease of notation that $\hat{T}_{(i)} = \hat{T}_i$
2: **for** *boot* in $1, \ldots,$ `BOOT` **do**
3:     Generate and iid bootstrap replicate $Z_{\pi,j}^{(l,boot)}(\mathbf{s})$, according to equation (19), for all $l$ and $j$
4:     Compute $\hat{T}_i^{boot}$, the estimate of $\hat{T}_i$ in bootstrap replicate *boot*, for $i \in \{1, \ldots, \eta\}$
5: **for** $u'$ in $1, \ldots, \eta$ **do**
6:     $\text{wFDR}^{boot}(u') \leftarrow \frac{\sum_{i=1}^{u'} a_i^{boot} T_i^{boot}}{\sum_{i=1}^{u'} a_i^{boot}}$
7:     $\text{wFDX}(u') \leftarrow \frac{1}{\texttt{BOOT}} \sum_{boot=1}^{\texttt{BOOT}} \mathbb{1}[\text{wFDR}^{boot}(u') > t]$
8: Compute $u = \text{argmax}_{u' \in 1,\ldots,\eta} \Big\{ \text{wFDX}(u) \leq \alpha \Big\}$
9: Reject hypotheses at the windows corresponding to statistics $\hat{T}_1, \ldots, \hat{T}_u$

Algorithm 2 was already proposed by Sun et al. (2015)[27], while Algorithm 3 is a novel construction. We have the following theorem about Algorithm 3.

**Theorem 1.** *Algorithm 3 achieves* asymptotic $(t, \alpha)-$*control of the frequentist wFDX.*

To our knowledge, the wFDX has been discussed only in the frequentist setting. Perone Pacifico et al. (2004) estimated this quantity by first identifying a confidence superset of the true-null regions. That is, given $S_{TN}$ as the true regions corresponding to the null hypothesis, they propose identifying a region $U$ such that $P(U \supset S_{TN}) \geq 1 - \alpha$. This region is used to estimate error-controlling quantities. However, though this method improves in power over a naive application of the Benjamini-Hochberg procedure[40], the confidence superset used is a conservative estimate, rendering this approach overly conservative[27,41,42]. Moreover, for our purposes, it does not naturally allow the incorporation of uncertainty in the background (i.e., $\mu(\mathbf{s})$). In the Bayesian setting, alternatively, Sun et al. (2015)[27] discussed how such testing quantities relate to simple classification problems. We adapt those ideas into Algorithm 3 for intuitive estimation of wFDX. By capturing the joint posterior distribution of the data and the mean function (equations (20) and (21)) we are able to test hypotheses with sharper error control and improved power.

### S2 Proofs

#### S2.1 Proof of Theorem 1

*Proof.* The frequentist wFDX at threshold $t$ is

$$\begin{aligned}
\text{wFDX}_t &= P\Bigg\{E\Bigg[\frac{\sum_{i=1}^{\eta} a_i(1-\theta_i)\delta_i}{\sum_{i=1}^{\eta} a_i\delta_i}\Bigg] > t\Bigg\} \\
&= P\Bigg\{E\Bigg[E\Big(\frac{\sum_{i=1}^{\eta} a_i(1-\theta_i)\delta_i}{\sum_{i=1}^{\eta} a_i\delta_i} \mid \mathbf{Z}\Big)\Bigg] > t\Bigg\} && \text{(iterated expectation theorem)} \\
&= P\Bigg\{E\Bigg[\frac{\sum_{i=1}^{\eta} a_i\delta_i E\Big((1-\theta_i) \mid \mathbf{Z}\Big)}{\sum_{i=1}^{\eta} a_i\delta_i}\Bigg] > t\Bigg\} \\
&= P\Bigg\{\frac{\sum_{i=1}^{\eta} a_i\delta_i P(\theta_i = 0 \mid \mathbf{Z}\Big)}{\sum_{i=1}^{\eta} a_i\delta_i} > t\Bigg\}
\end{aligned}$$

The quantity $P(\theta(s_i) = 0 \mid \mathbf{Z})$ is approximated as in equation (21). Let $R_t$ be an assumed set of rejected windows. Then,

$$\begin{aligned}
\widehat{\text{wFDX}}_t &= P\Bigg\{\frac{\sum_{i=1}^{\eta} a_i\hat{T}_i \cdot \mathbb{1}(s_i \in R_t)}{\sum_{i=1}^{\eta} a_i \cdot \mathbb{1}(s_i \in R_t)} > t\Bigg\} \\
&= \frac{1}{\texttt{BOOT}} \sum_{boot=1}^{\texttt{BOOT}} \mathbb{1}\Bigg\{\frac{\sum_{i=1}^{\eta} a_i\hat{T}_i^{boot} \cdot \mathbb{1}(s_i \in R_t)}{\sum_{i=1}^{\eta} a_i \cdot \mathbb{1}(s_i \in R_t)} > t\Bigg\}
\end{aligned}$$

where the final equality follows from the fact that the bootstrap replicates are all iid by construction. This term is always less than or equal to $\alpha$ by step 8 of Algorithm 3. Therefore, control of the estimated Bayesian $\widehat{\text{wFDX}}_t$ implies control of the frequentist $\text{wFDX}_t$ asymptotically in the number of windows. □

##### S2.2 Proof of Proposition 1

To prove Proposition 1, we first introduce some notation. Let $\text{FDR}_\lambda = E[\text{FDP}_\lambda]$, where $\text{FDP}_\lambda$ is the proportion of rejected area in 2D that does not overlap with truly non-null area in 2D. Recall that $\delta(s_w) \in \{0, 1\}$ is the decision made at location $s_w$. For a decision made based on a process derived from scanning windows of size $B$, $\delta(s_w) = 1$ implies a significant difference in a triangular region on the Hi-C contact matrix. This triangular region is an isosceles right triangle with hypotenuse of length $B$, and this hypotenuse begins at location $s_w$ and ends at location $s_w + B - 1$. This triangle, denoted $\Delta_\delta^B(s_w)$, has area $B^2/4$. We similarly define $\theta^{B'}(s_{w'}) = 1$ as implying a truly differential triangular region $\Delta_\theta^{B'}(s_{w'})$, with length $B'$ hypotenuse beginning at location $s_{w'}$. Then, $\delta(s_w)$ is considered a false discovery if $\delta(s_w) = 1$ and there does not exist some pair $(s_{w'}, B')$ such that $\theta^{B'}(s_{w'}) = 1$, $s_{w'} \leq s_w$, and $B' \geq B + s_w - s_{w'}$. Briefly, $\delta(s_w) = 1$ is a false discovery if the corresponding triangular region $\Delta_\delta^B(s_w)$ is not completely covered by some other triangular region $\Delta_\theta^{B'}(s_{w'})$ corresponding to a truly differential region.

Define $S_{FP}$ as the false positive area and $R$ as the rejection area, and let $\nu(\cdot)$ be the counting measure on $S$. Then for some small constant $c_0 \in (0, 1)$, control of the FDR on the 1D NNGP by definition controls the following quantity:

$$\begin{aligned}
\text{FDR} &= E\{\text{FDP}\} \\
&= E\Big\{\frac{\nu(S_{FP})}{\nu(R)}\mathbb{1}(\nu(R) > c_0)\Big\} \\
&= E\left\{\frac{\sum_{w=1}^{W}\Big[\delta(s_w)\cdot\mathbb{1}(\nexists\,(s_{w'},\, B') \ni \theta^{B'}(s_{w'}) = 1,\, s_{w'} \leq s_w,\, B' \geq B + s_w - s_{w'})\Big]}{\sum_{w=1}^{W}\delta(s_w)}\mathbb{1}(\sum_{w=1}^{W}\delta(s_w) > c_0)\right\} \\
&\leq \alpha.
\end{aligned} \tag{22}$$

Now, let $\lambda(\cdot)$ be the Lebesgue measure on $S \times S$, such that $\lambda(\cdot)$ measures the size of the area covered by $\Delta_\delta^B(s)$ and $\Delta_\theta^{B'}(s)$ for all $s \in S$. That is, if one rejection is made in a window of size $B$, then $\lambda(R) = B^2/4$, but if $m$ rejections are made, then $\lambda(R)$ captures the area of the overlap of all rejections, that is

$$\lambda(R) = \lambda\Bigg(\bigcup_{w=1}^{W}\Big[\mathbb{1}(\delta(s_w) = 1)\cdot\Delta_\delta^B(s_w)\Big]\Bigg). \tag{23}$$

Thus, as in equation (22), we can similarly define $\text{FDR}_\lambda$, which measures the false discoveries based on overlaps among rejections with truly differential regions as

$$
\begin{aligned}
\text{FDR}_\lambda &= E\{\text{FDP}_\lambda\} \\
&= E\{\frac{\lambda(S_{FP})}{\lambda(R)} \mathbb{1}(\lambda(R) > c_0)\} \\
&= E\left\{ \frac{\lambda\Big(\bigcup_{w=1}^{W} \Big[\mathbb{1}(\delta(s_w) = 1) \cdot \Delta_\delta^B(s_w)\Big]\Big) - \lambda\Big(\bigcup_{w=1}^{W} \Big[\mathbb{1}(\delta(s_w) = 1) \cdot \Delta_\delta^B(s_w)\Big] \cap \Delta_{\theta=1}\Big)}{\lambda\Big(\bigcup_{w=1}^{W} \Big[\mathbb{1}(\delta(s_w) = 1) \cdot \Delta_\delta^B(s_w)\Big]\Big)} \mathbb{1}(\lambda(R) > c_0) \right\}
\end{aligned} \tag{24}
$$

where the region

$$
\Delta_{\theta=i} = \bigcup_{s \in S} \bigcup_{B \in \mathcal{B}} \Big[\Delta_\theta^B \cdot \mathbb{1}(\theta^B(s) = i)\Big] \tag{25}
$$

where $\mathcal{B}$ is the set of all possibly hypotenuse lengths for truly differential triangular regions. Now we can make a statement about false discovery control based on overlap of differential regions on the Hi-C contact map.

*Proof.* We only need to show that $\text{FDP}_\lambda \leq \text{FDP}$ since $x \leq y \Rightarrow E[x] \leq E[y]$. When the total number of rejections $m = 0$, $\text{FDP}_\lambda = \text{FDP} = 0 \leq \alpha$, so the inequality is trivially true. When $m = 1$, $\nu(S_{FP}) = 0 \Rightarrow \lambda(S_{FP}) = 0$, such that again $\text{FDP}_\lambda = \text{FDP} = 0 \leq \alpha$. However, if $\nu(S_{FP}) = 1$, then $\text{FDP} = 1$. But, letting $s$ be the start of the rejected region, $\text{FDP}_\lambda = \frac{B^2/4 - \lambda\left(\Delta_\delta^B(s) \cap \Delta_{\theta=1}\right)}{B^2/4}$, where the second term in the numerator is in $[0, B^2/4)$, such that $\text{FDP}_\lambda \leq \text{FDP} \leq \alpha$.

The inductive hypothesis is that for $1 \leq m < W$ rejections, $\text{FDP}_\lambda \leq \text{FDP}$. We need to show that this implies that for $m+1$ rejections, the inequality still holds. First, assume we can order the regions $s_1 \ldots, s_W$ such that the first $m$ regions $s_1, \ldots, s_m$ correspond to the first $m$ rejections, and that $s_{m+1}$ is the $(m+1)^{th}$ rejection. Then,

$$
\begin{aligned}
\text{FDP}_\lambda &= \frac{\lambda\Big(\bigcup_{w=1}^{m} \Delta_\delta^B(s_w) \cup \Delta_\delta^B(s_{m+1})\Big) - \lambda\Big(\Big\{\bigcup_{w=1}^{m} \Delta_\delta^B(s_w) \cup \Delta_\delta^B(s_{m+1})\Big\} \cap \Delta_{\theta=1}\Big)}{\lambda\Big(\bigcup_{w=1}^{m} \Delta_\delta^B(s_w) \cup \Delta_\delta^B(s_{m+1})\Big)} \\
&= \frac{\lambda\Big(\bigcup_{w=1}^{m} \Delta_\delta^B(s_w) \cap \Delta_{\theta=0}\Big) + \lambda\Big(\Delta_\delta^B(s_{m+1}) \cap \Delta_{\theta=0}\Big) - \lambda\Big(\bigcup_{w=1}^{m} \Delta_\delta^B(s_w) \cap \Delta_\delta^B(s_{m+1}) \cap \Delta_{\theta=0}\Big)}{\lambda\Big(\bigcup_{w=1}^{m} \Delta_\delta^B(s_w)\Big) + B^2/4 - \lambda\Big(\bigcup_{w=1}^{m} \Delta_\delta^B(s_w) \cap \Delta_\delta^B(s_{m+1})\Big)} \\
&\leq \frac{\lambda\Big(\bigcup_{w=1}^{m} \Delta_\delta^B(s_w) \cap \Delta_{\theta=0}\Big) + \lambda\Big(\Delta_\delta^B(s_{m+1}) \cap \Delta_{\theta=0}\Big)}{\lambda\Big(\bigcup_{w=1}^{m} \Delta_\delta^B(s_w)\Big) + B^2/4}
\end{aligned} \tag{26}
$$

Where the final inequality comes from assuming that that $\Delta_\delta^B(s_{m+1})$ is completely disjoint from $\bigcup_{w=1}^{m} \Delta_\delta^B(s_w)$. Then, the second term in the numerator is 0 if the $(m+1)^{th}$ discovery is true, and is in the interval $(0, B^2/4]$ otherwise. In the former case,

$$\frac{\lambda\Big(\bigcup_{w=1}^m \Delta_\delta^B(s_w)\cap\Delta_{\theta=0}\Big)}{\lambda\Big(\bigcup_{w=1}^m \Delta_\delta^B(s_w)\Big)+B^2/4} < \frac{\lambda\Big(\bigcup_{w=1}^m \Delta_\delta^B(s_w)\cap\Delta_{\theta=0}\Big)}{\lambda\Big(\bigcup_{w=1}^m \Delta_\delta^B(s_w)\Big)} \le \mathrm{FDP} \qquad \text{(inductive hypothesis)}$$

.

For the latter case,

$$\begin{aligned}
\mathrm{FDP}_\lambda &= \frac{\lambda\Big(\bigcup_{w=1}^m \Delta_\delta^B(s_w)\cap\Delta_{\theta=0}\Big)+\lambda\Big(\Delta_\delta^B(s_{m+1})\cap\Delta_{\theta=0}\Big)}{\lambda\Big(\bigcup_{w=1}^m \Delta_\delta^B(s_w)\Big)+B^2/4}\\
&= \frac{\lambda\Big(\bigcup_{w=1}^m \Delta_\delta^B(s_w)\cap\Delta_{\theta=0}\Big)}{\lambda\Big(\bigcup_{w=1}^m \Delta_\delta^B(s_w)\Big)+B^2/4} + \frac{B^2/4-\lambda\Big(\Delta_\delta^B(s_{m+1})\cap\Delta_{\theta=1}\Big)}{\lambda\Big(\bigcup_{w=1}^m \Delta_\delta^B(s_w)\Big)+B^2/4}\\
&\le \frac{\lambda\Big(\bigcup_{w=1}^m \Delta_\delta^B(s_w)\cap\Delta_{\theta=0}\Big)}{\lambda\Big(\bigcup_{w=1}^m \Delta_\delta^B(s_w)\Big)+1} + \frac{B^2/4-\lambda\Big(\Delta_\delta^B(s_{m+1})\cap\Delta_{\theta=1}\Big)}{\lambda\Big(\bigcup_{w=1}^m \Delta_\delta^B(s_w)\Big)+B^2/4} \qquad (\mathrm{B}\ge 2)\\
&\le \frac{\sum_{w=1}^m \mathbb{1}(\nexists\,(s_{w'},\,B')\ni\theta^{B'}(s_{w'})=1,\ s_{w'}\le s_w,\ B'\ge B+s_w-s_{w'})}{m+1}+\frac{1}{m+1}\\
&\le \mathrm{FDP}
\end{aligned}$$

where the penultimate inequality comes from the inductive hypothesis, and that $A/B \le C/D \Rightarrow A/(B+1) \le C/(D+1)$ for $A, B, C, D > 0$. This concludes the proof by induction.

□

### S3 Polymer simulation details

#### S3.1 Polymer forces

The following potentials were used for the polymer simulations. Here and below distances are measured in units of monomer diameter, 20nm. Neighboring monomers were held together by harmonic bonds with a potential $U = 100(r-1)^2$ (here and below, all energies are in units of kT). Polymer *stiffness* is modeled with a three-point interaction term, with the potential $U = S(1-\cos(\alpha))$, where $\alpha$ is the angle between neighboring bonds, and $S$ is a stiffness parameter taken to be 1.5.

To allow chain passing, which represents activity of topoisomerase II, we used a soft-core potential for interactions between monomers, similar to the one used in Fudenberg et al. (2016)[46]. All monomers interacted via a repulsive potential

$$U = 4\Big\{-1+\Big[\frac{1.05r}{\sqrt{6/7}}\Big]^{12}\cdot\Big[\Big(\frac{1.05r}{\sqrt{6/7}}\Big)^2-1\Big]\cdot\frac{823543}{46656}\Big\}. \qquad (27)$$

This is a fast and efficient potential designed to be a constant of 4 kT up to $r = 0.7-0.8$, and then quickly go to 0 at $r = 1.05$ together with its first derivative. Simulations were performed in periodic boundary conditions, with the size of the cubic box set to achieve spatial *density* of 0.1.

We used Langevin integrator with error tolerance (described in the OpenMM user manual) of 0.01, and with the collision rate of 0.003 as defined in the polychrom library. For the sake of computational efficiency, we simulated 4 copies of the system together in one simulation. This was done because performance of OpenMM with 60,000 monomers is only about 2 times slower than with 15,000 monomers, thus yielding $\sim 4 \cdot 0.5 = 2\times$ more results per unit time.

##### S3.2 Loop extrusion simulations

Loop extrusion dynamics were simulated as in Fudenberg et al. (2016)[46] with minor modifications. To simulate loop extrusion, we performed one-dimensional simulation of Loop Extruding Factors (LEFs) on a binary lattice with one monomer representing one polymer monomer, or 2 kb. In interphase cells of mammals, cohesin acts as a LEF, with CTCF acting as extrusion barriers.

For loop extrusion simulations, we used *processivity* of 280 kb and *separation* of 280 kb. Processivity and separation are as defined in Fudenberg et al. (2016): processivity is the average size of a loop that would be extruded by a LEF without obstacles, formally defined as

$$\text{processivity (kb)} = 2 \cdot \frac{1}{\text{dissociation rate of a LEF}} \cdot \frac{\text{kb}}{\text{monomer}}$$

where 2 comes in because both legs of a LEF translocate by 1 monomer during one step of loop extrusion, thus increasing the loop size by 2 monomers. Separation is defined as the inverse density of LEFs:

$$\text{separation (kb)} = \frac{\#\text{monomers}}{\#\text{LEFs}} \cdot \frac{\text{kb}}{\text{monomer}}.$$

At each step, two legs of each loop extruder attempt to translocate by 1 monomer, away from each other. If a leg encounters an occupied site, it waits until the site becomes free. LEFs in our simulations are *two-sided*, meaning that if one leg is stalled, another leg continues to translocate. LEFs have a constant dissociation rate defined by processivity, the dissociation rate is independent of whether any legs are stalled. For the sake of computational efficiency, we simulate a fixed number of LEFs: once a LEF dissociates, it re-associates in a random unoccupied location.

We simulate TAD boundaries (CTCF sites) as in Fudenberg et al. (2016). We simulate boundaries as bi-directional or uni-directional (i.e. able to stop LEFs travelling both directions, or stop LEFs travelling in one direction only) depending on a simulation performed. Each boundary has a certain probability to capture a translocating LEF. Once captured, a leg of a LEF remains immobile forever, i.e. until the LEF dissociates.

##### S3.3 Coupling loop extrusion and polymer simulations

As previously, we simulate LEFs as harmonic polymer bonds connecting two monomers. The energy of the bond was chosen to be softer than harmonic polymer bonds: $U = 25(r - 0.5)^2$. To simulate translocation of a LEF, a bond was moved to the neighboring monomer(s) according to the LEF dynamics. Softer bonds were necessary to avoid excessive jerking of the system when the bond is re-assigned. We performed 600 steps of Langevin dynamics between adjusting positions of LEF bonds.

#### S4 Supplementary figures

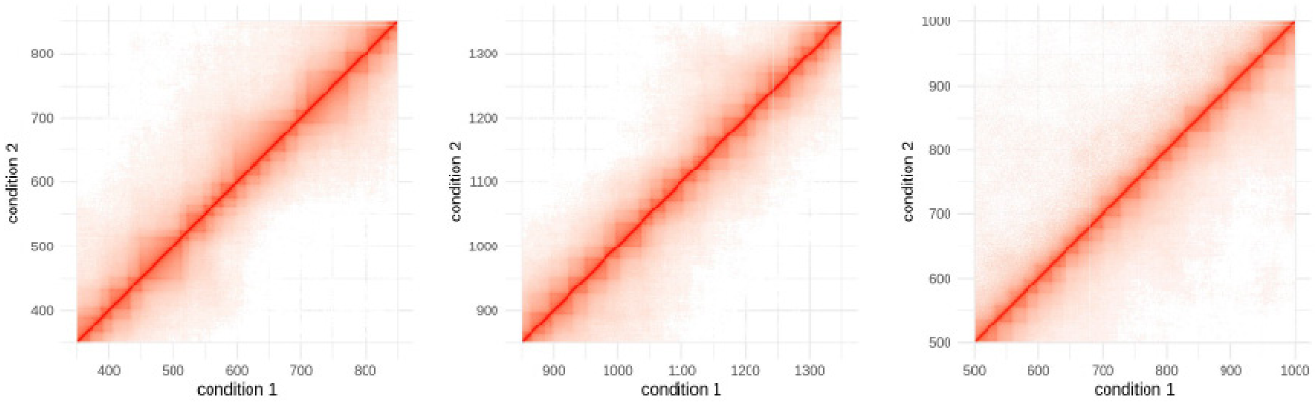


Figure S1: Visualization of example replicates from condition 1 versus condition 2 in (from left to right) simulations 1 through 3.

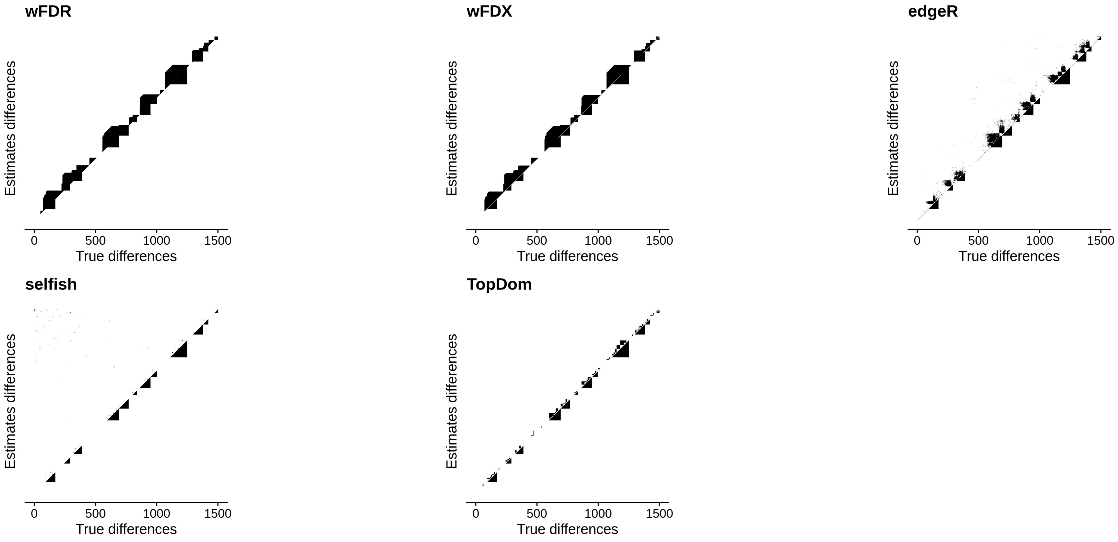


Figure S2: True differences vs. estimated differences at confidence level $\alpha = 0.01$ across $W = 1,500$ locations for 5 methods in simulation 1 with contact radius 6. The sixth method, multiHiCcompare, did not run on this dataset.

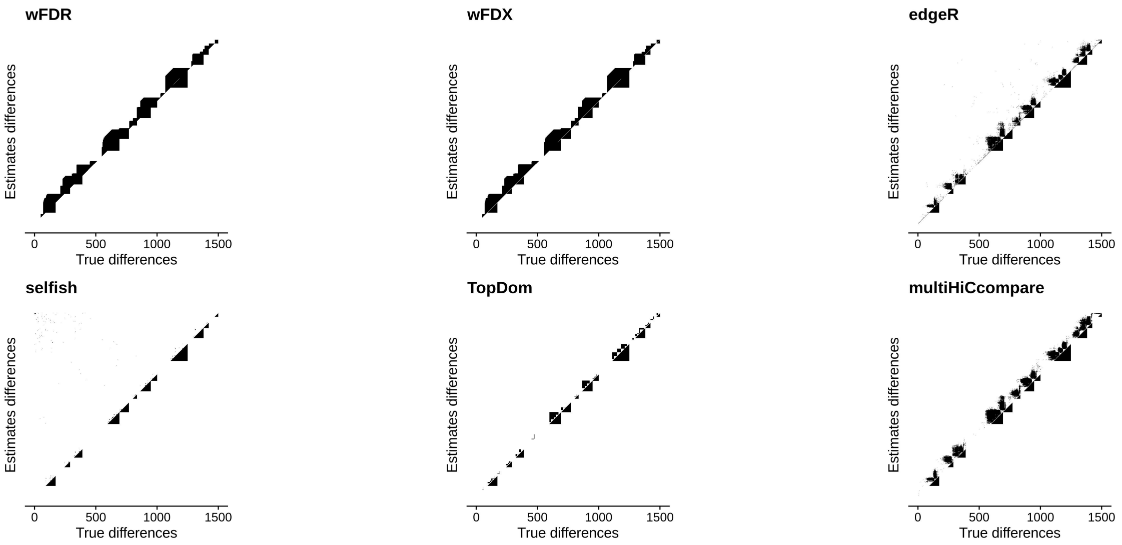


Figure S3: True differences vs. estimated differences at confidence level $\alpha = 0.01$ across $W = 1,500$ locations for 6 methods in simulation 1 with contact radius 8.

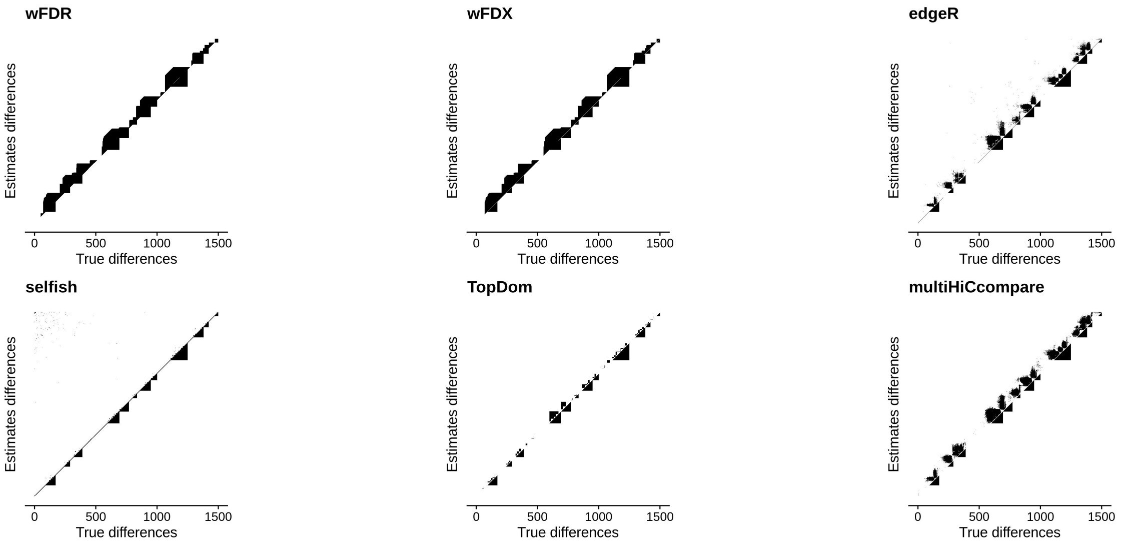


Figure S4: True differences vs. estimated differences at confidence level $\alpha = 0.01$ across $W = 1,500$ locations for 6 methods in simulation 1 with contact radius 10.

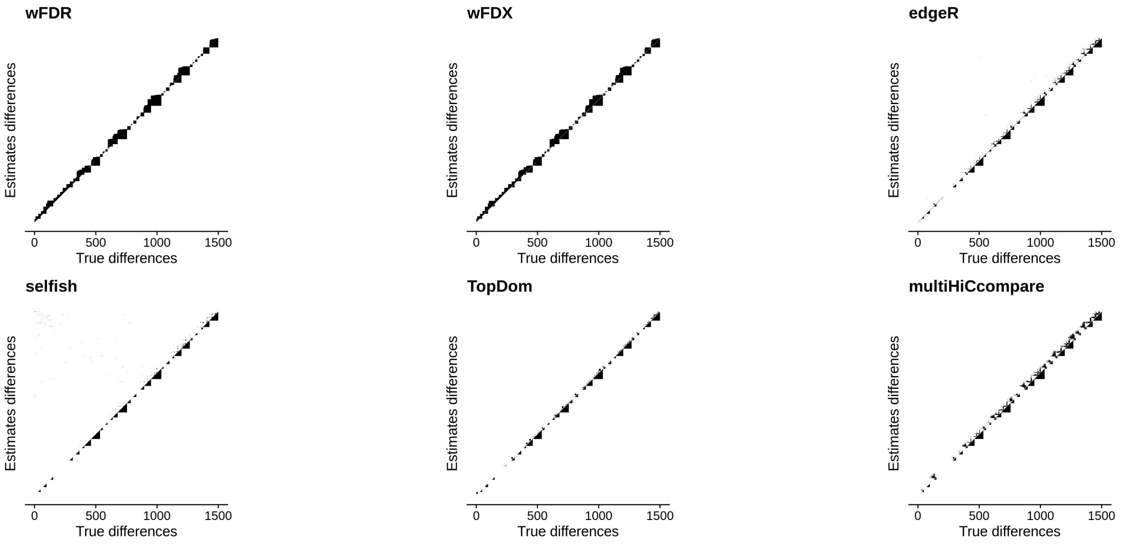

Figure S5: True differences vs. estimated differences at confidence level $\alpha = 0.01$ across $W = 1,500$ locations for 6 methods in simulation 2 with contact radius 6.

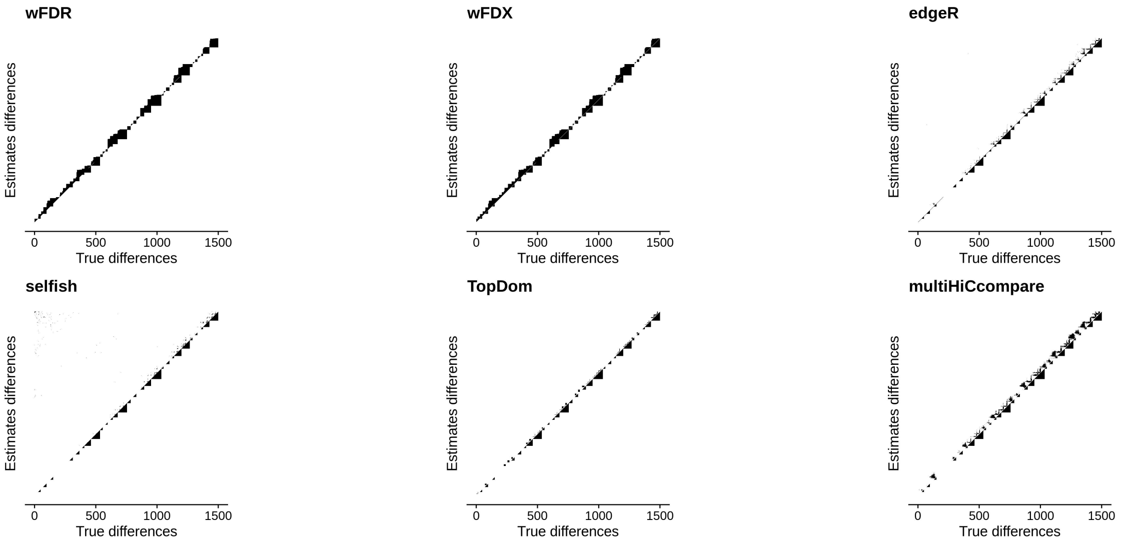

Figure S6: True differences vs. estimated differences at confidence level $\alpha = 0.01$ across $W = 1,500$ locations for 6 methods in simulation 2 with contact radius 8.

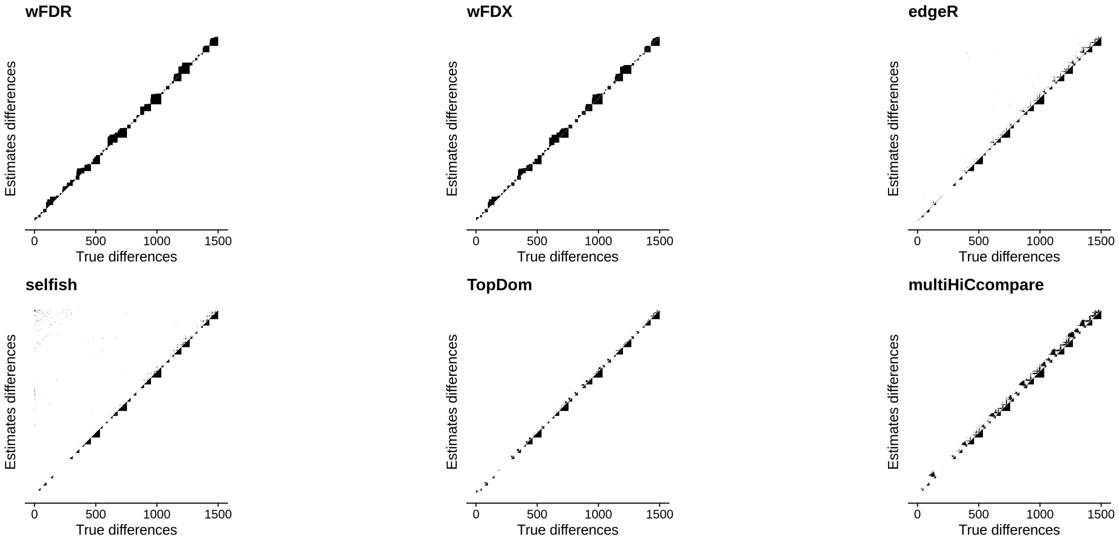


Figure S7: True differences vs. estimated differences at confidence level $\alpha = 0.01$ across $W = 1,500$ locations for 6 methods in simulation 2 with contact radius 10.

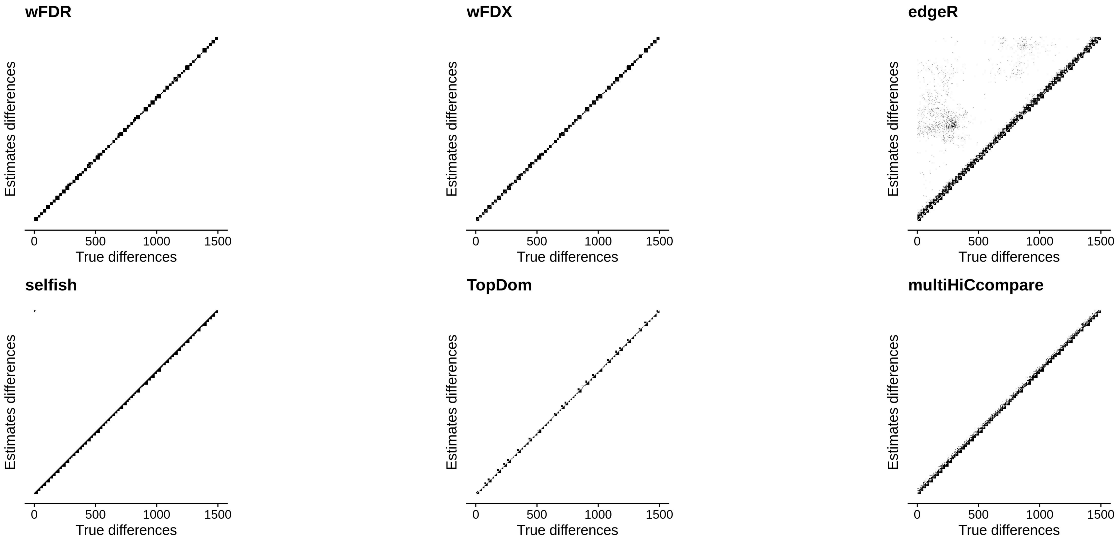


Figure S8: True differences vs. estimated differences at confidence level $\alpha = 0.01$ across $W = 1,500$ locations for 6 methods in simulation 3 with contact radius 6.

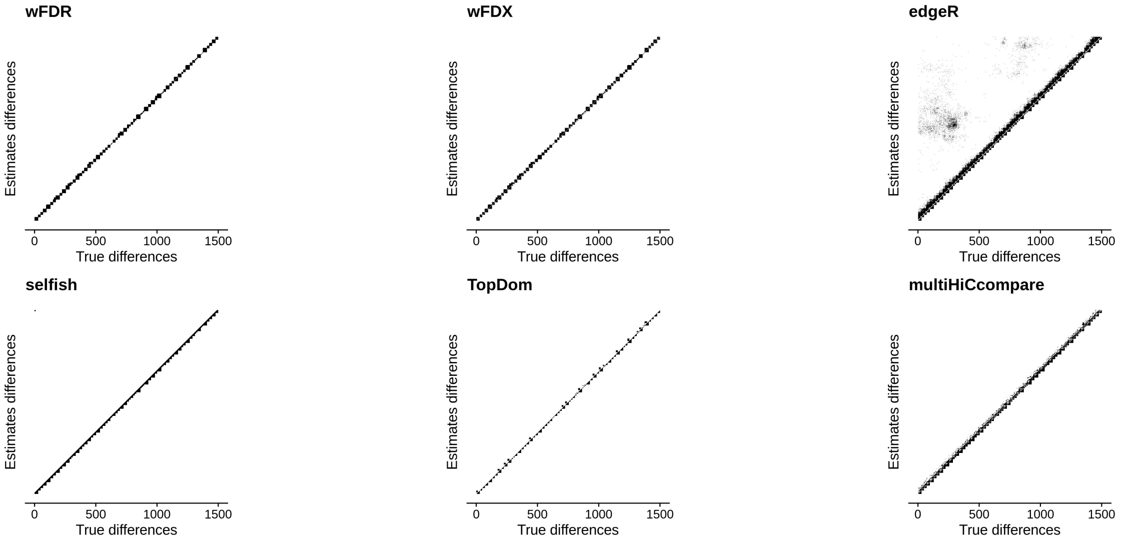


Figure S9: True differences vs. estimated differences at confidence level $\alpha = 0.01$ across $W = 1,500$ locations for 6 methods in simulation 3 with contact radius 8.

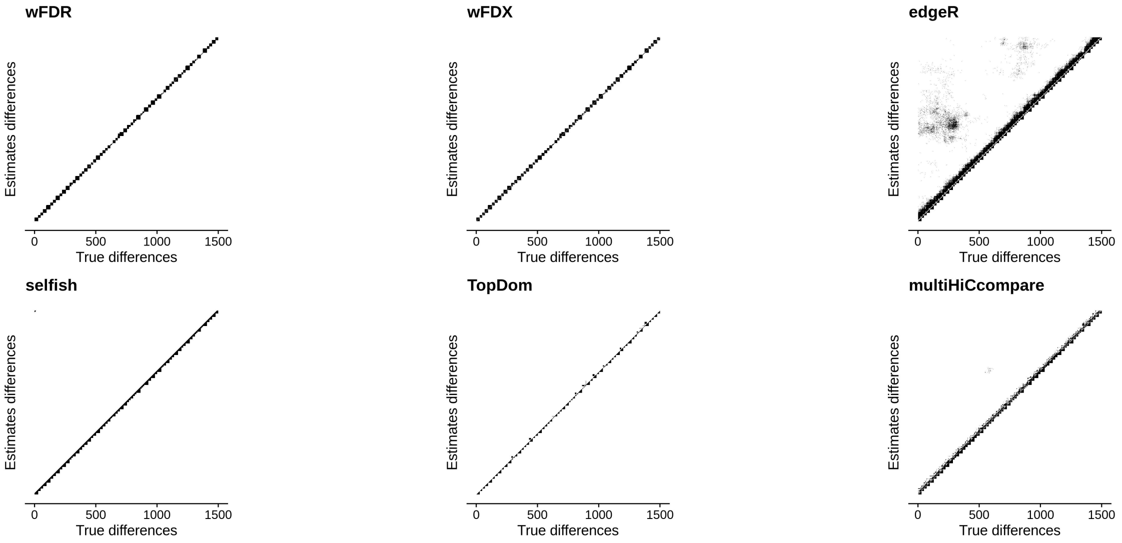


Figure S10: True differences vs. estimated differences at confidence level $\alpha = 0.01$ across $W = 1,500$ locations for 6 methods in simulation 3 with contact radius 10.

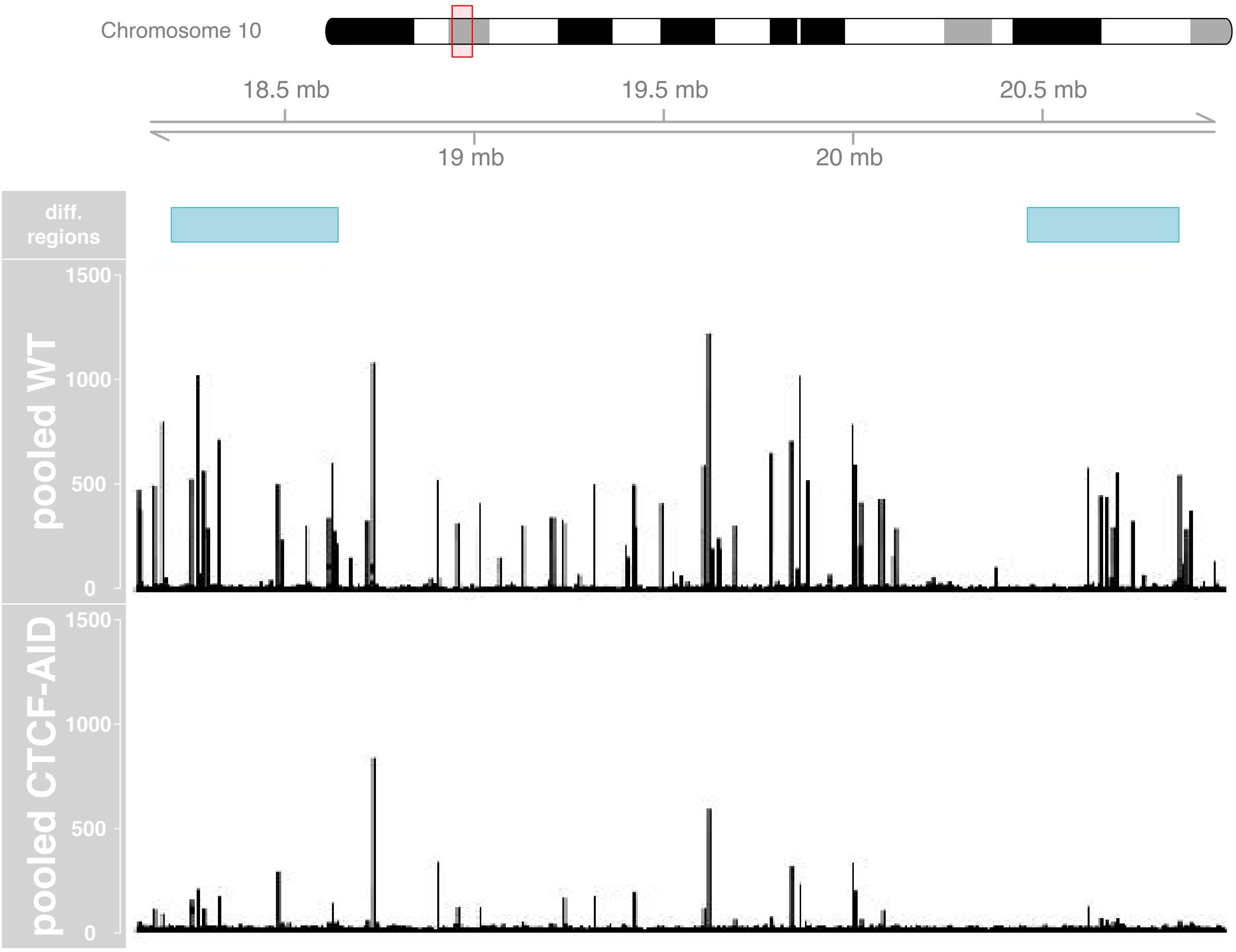


Figure S11: Sample browser shot of significantly differential regions in the CTCF-AID analysis.

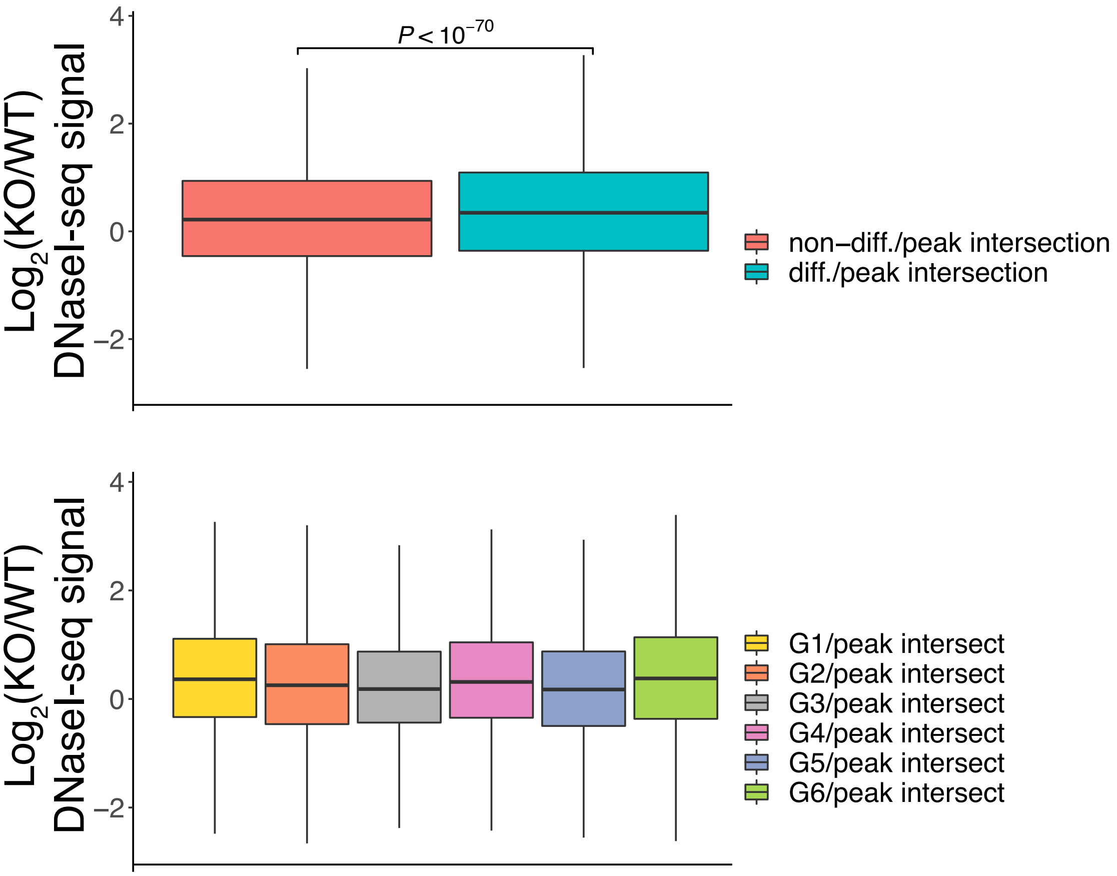


Figure S12: $\text{Log}_2$ fold change between wild type (WT) and Chd4-knockout (KO) of the average DNaseI-seq signal within DNaseI-seq peak regions. Peaks, originally called on these data by Goodman et al.[28], were intersected with locdiffr's differential and non-differential regions, as well as groups $G1 - G6$ as described in Fig. 4c. Unlike the boxplots for DNaseI-seq in Fig. 4a–b, these boxplots all show a positive median $\log_2$ ratio. This is due to systematically lower DNaseI-seq peak signal, but systematically higher background DNaseI-seq signal, in the WT experiments when compared against KO (see also supplementary Fig. S13). Despite this change, the same trends hold: locdiffr differential regions are enriched for increased accessibility, and when intersected with peak calling information from HiCCUPS, groups $G1$, $G4$, and $G6$ show the largest gain in accessibility in the KO experiment over the WT experiment. There were 59,352 peaks in non-differential regions, and 61,696 in differential regions. The $P$-value was computed using a two-tailed Wilcoxon rank-sum test. Outliers were omitted from figure for ease of visualization.

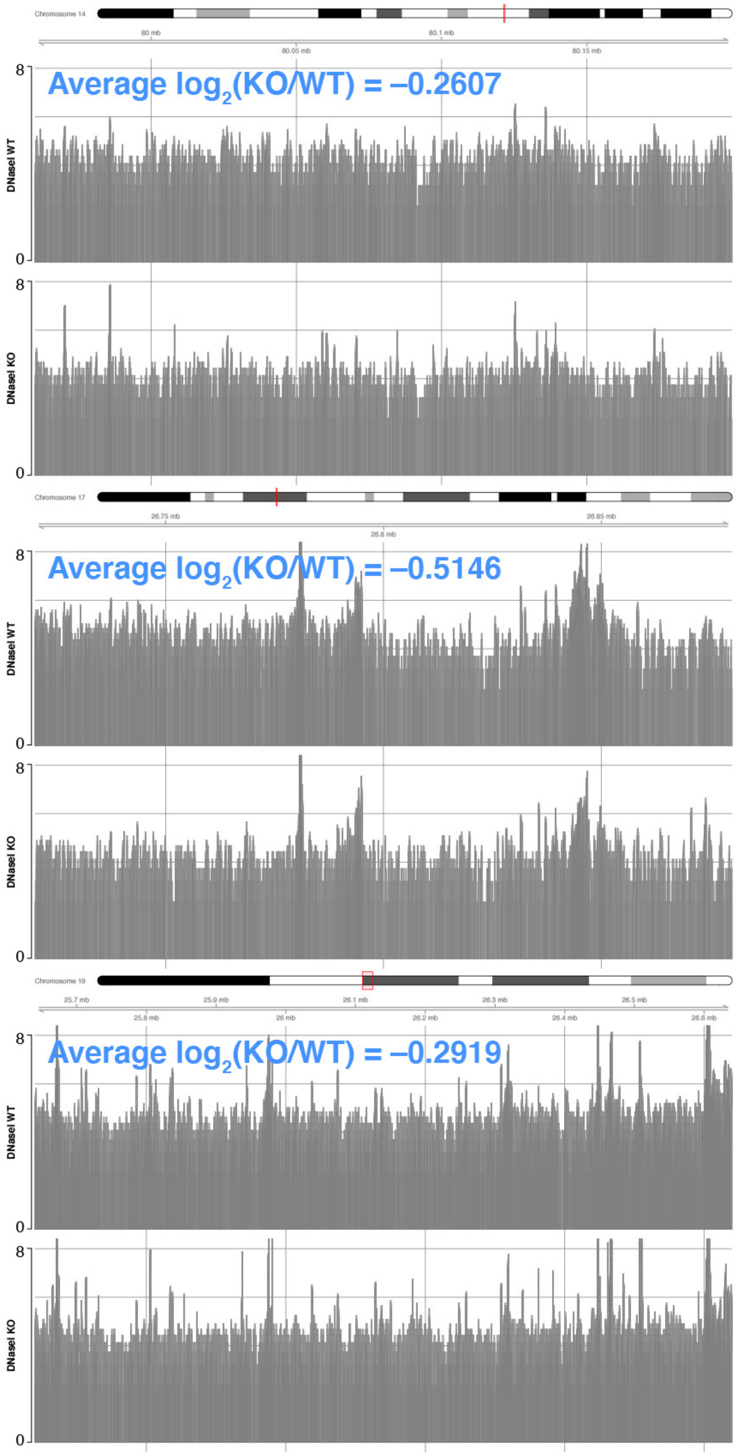


Figure S13: Comparison of average background signal in wild type (WT) and Chd4-knockout (KO) cells in 3 example background regions. DNaseI-seq signal is shown on log scale.

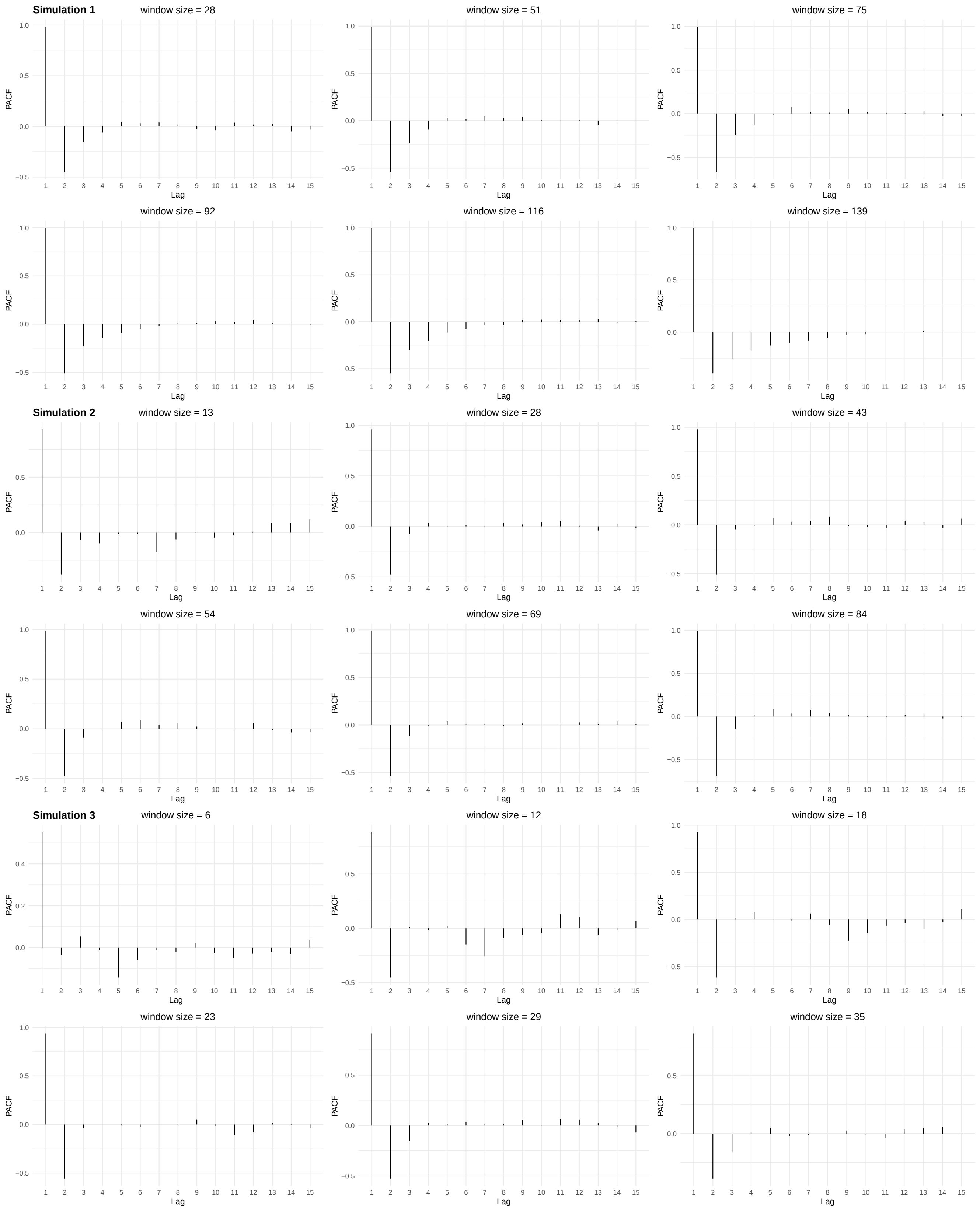


Figure S14: Partial autocorrelation functions for several window sizes after subtracting the estimated mean trend for the all simulations with contact radius 8.

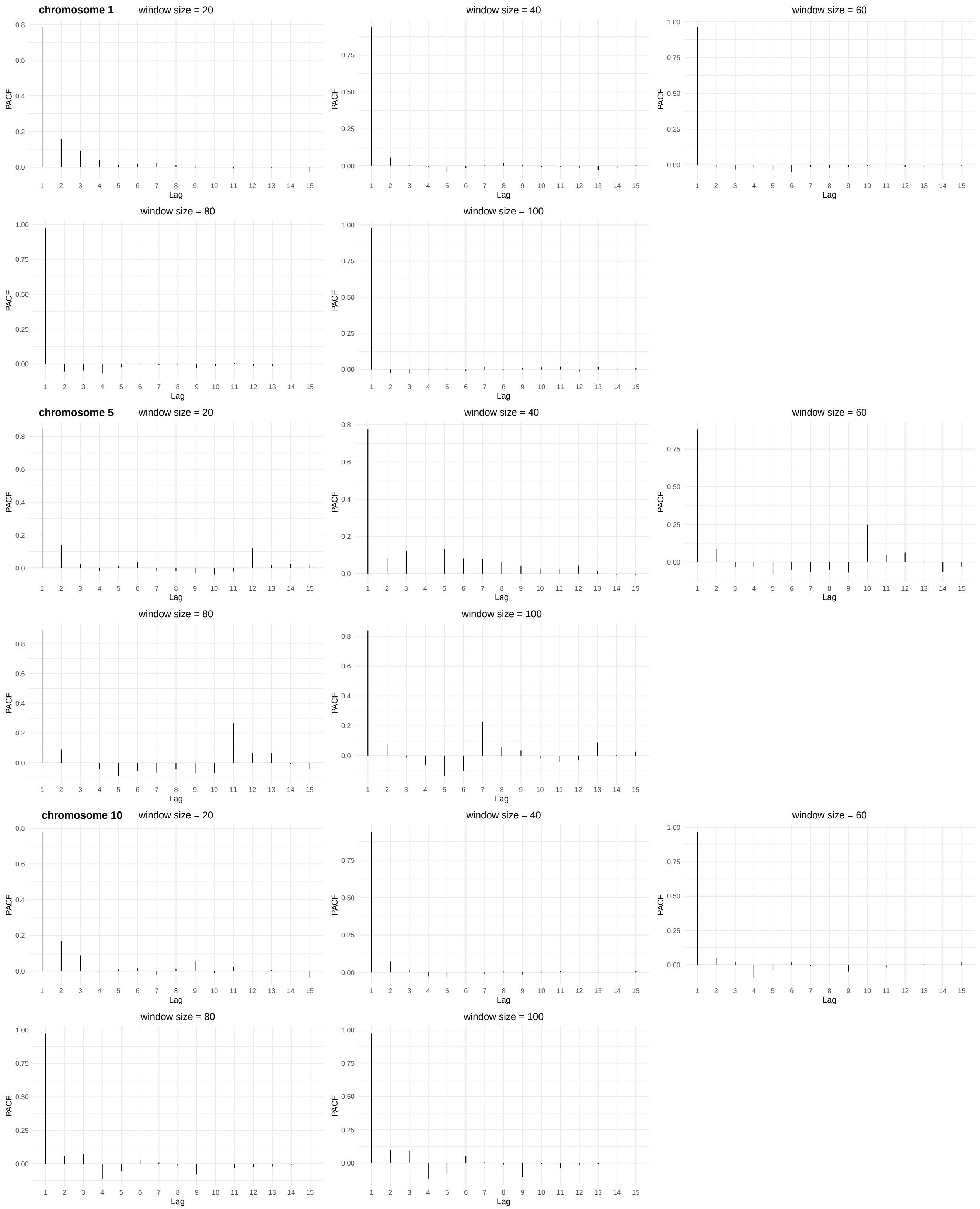


Figure S15: Partial autocorrelation functions for several chromosome/window size pairs after subtracting the estimated mean trend for the CTCF-AID analysis.

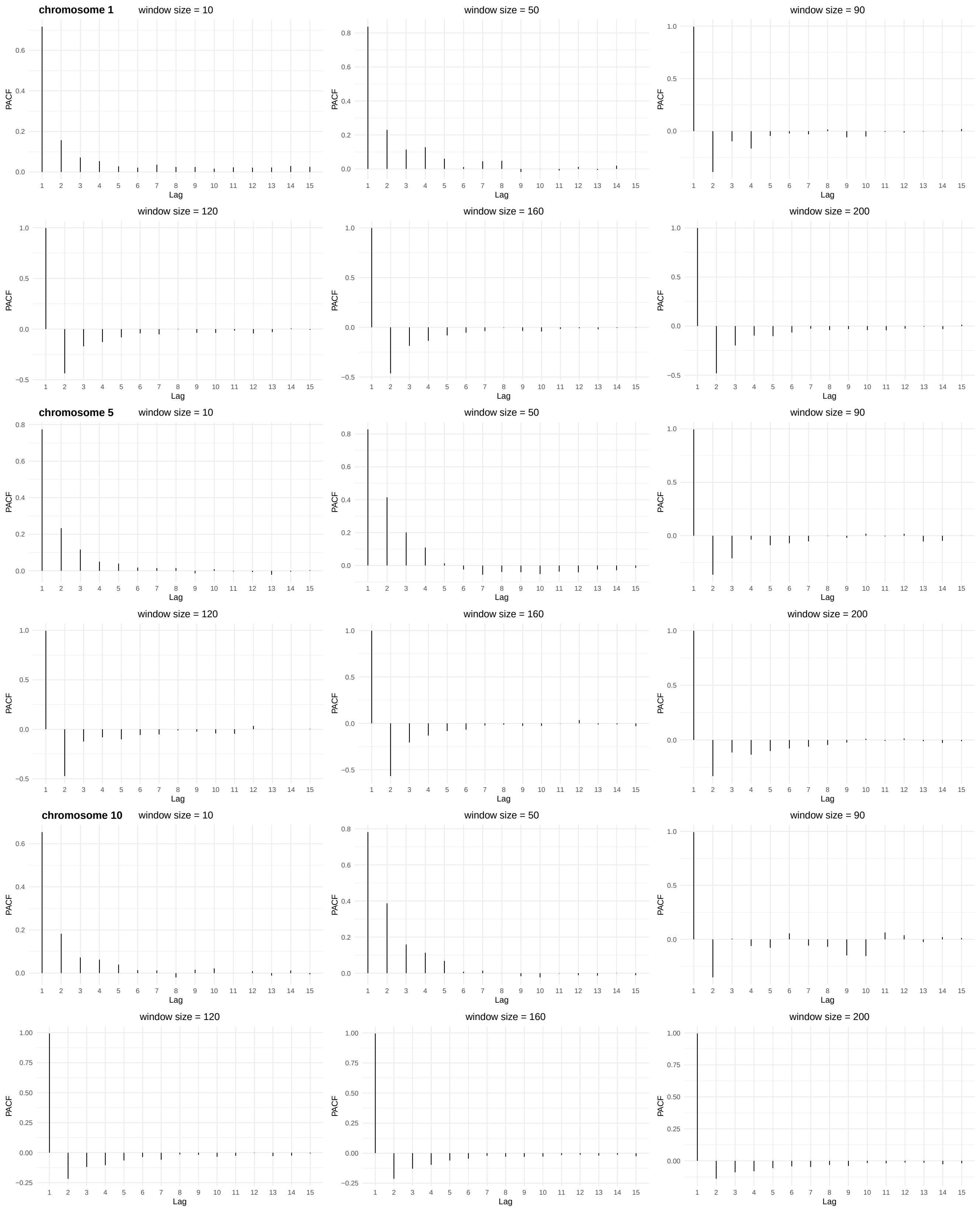


Figure S16: Partial autocorrelation functions for several chromosome/window size pairs after subtracting the estimated mean trend for the Chd4-knockout analysis.
